# Mapping early patterning events in human neural development using an *in-vitro* microfluidic stem cell model

**DOI:** 10.1101/2025.05.23.655442

**Authors:** Gaurav Singh Rathore, Matias Ankjær, Pedro Rifes, Erno Hänninen, Fátima Sanchís Calleja, Charlotte Rusimbi, Janko Kajtez, Amalie Holm Nygaard, Louise Saviø Piilgaard, Ugne Dubonyte, Zehra Abay-Nørgaard, Jens Bager Christensen, Kristoffer Lihme Egerod, J. Gray Camp, Tune H Pers, Barbara Treutlein, Agnete Kirkeby

## Abstract

Stem cell models can provide insights into human brain development at embryonic stages which are normally inaccessible. We previously developed the Microfluidic Stem Cell Regionalisation (MiSTR) model, which recapitulates the rostro-caudal patterning of human neural tube through a WNT activation (WNTa) gradient. Through temporal single cell transcriptomics of rostro-caudal and dorso-ventral gradient-patterned MiSTR, we found that rostro-caudal subtypes were regionally specified and fate-determined already during the late epiblast stage, several days before onset of neuralisation at day 3-4. Rostral cells were characterised by expression of HESX1 and SHISA2 during pre-neuralisation and PAX6 during early neuralisation, whereas caudal cells expressed FST and HOXA1 during pre-neuralisation and SOX1 as the dominant neuralising factor. In contrast to the early rostro-caudal specification, response to ventralisation in telencephalic progenitors was developmentally delayed and occurred around day 9. We further uncovered temporal events in human midbrain-hindbrain boundary formation and ventral forebrain patterning, contributing new knowledge on early human neural region-specification.

## Introduction

One of the major challenges in translational applications of stem cell therapies is the incomplete knowledge of lineage relationships during early development, particularly in the brain, which possesses a very high degree of cellular diversity and complex tissue organisation. Although single cell RNA sequencing (scRNAseq) technologies have been instrumental in producing detailed and comprehensive cellular atlases of mouse and human embryonic development, few datasets are currently available at the very earliest fetal stages around 2-5 post-conception week (PCW) (La Manno et al. 2021, Braun et al. 2023, Mannens et al. 2024). Those few human datasets available from these stages only cover a total of a few thousand cells from the presumptive neural tube (Xu et al. 2023, Zeng et al. 2023), which limits confidence in identification of subregionalised cell types and temporal expression profiles at this highly dynamic time window of development.

Organoid models have proven to be excellent models for studying the developmental basis of early organ patterning and tissue organisation (Lancaster et al. 2013, Bagley et al. 2017, Quadrato et al. 2017, Xiang et al. 2017, Atamian et al. 2024). However, while organoids can self-organize into specific subregions of the brain, they offer limited surrogacy to the overall regionalisation of the neural tube forming the developing human brain. We have previously established a microfluidically controlled neurodevelopmental model, the Microfluidic Stem Cell Regionalisation (MiSTR) model, which recapitulates with reproducibility the rostro-caudal patterning of the developing neural tube spanning from forebrain to hindbrain (Rifes et al. 2020). MiSTR model relies on exposing a layer of pluripotent human embryonic stem cells (hESCs) to a gradient of the GSK3 inhibitor CHIR99021 (CHIR) to progressively activate WNT signalling, thereby inducing rostral-to-caudal neural regionalisation, mimicking early neural tube patterning (Kirkeby et al. 2012, Rifes et al. 2020). The model can be combined with the addition of SHH to mimic either the dorsal or the ventral axis of the neural tube (Rifes et al. 2020).

Other microfluidic and organoid-based stem cell models have applied the same concept of a CHIR-induced WNT activation (WNTa) gradient to successfully induce rostro-caudal patterning of neural cells from forebrain to hindbrain with or without the addition of SHH (Scuderi et al. 2024, Xue et al. 2024). These studies collectively confirm across several labs that a simple combination of a WNTa gradient in the presence or absence of SHH signalling robustly recapitulates rostro-caudal and dorso-ventral axial neural tube development in vitro. While uNTL (Xue et al. 2024), and Duo-MAPS (Scuderi et al. 2024) apply perpendicular gradients of WNT and SHH signalling, the MiSTR model is based on a single tightly controlled linear morphogen gradient generated by precision pumps (Rifes et al. 2020). Single-cell transcriptomic datasets from these human-specific engineered models of neural tube patterning can help bridge the gap in our understanding of the earliest stages of human neural development by providing access to hundreds of thousands of cells representing the earliest stages of regional neural patterning. In this study, we performed extensive temporal scRNAseq of rostro-caudal fate development in the MiSTR model, across both the dorsal and the ventral neural tube. In addition to previously published rostro-caudal models (Rifes et al. 2020), we also include a third gradient model to capture the dorso-ventral regionalization of the forebrain, recapitulating the formation of the ganglionic eminences and the dorsal telencephalon. This allowed us to generate a comprehensive temporal trajectory map of early neural tube patterning along both the rostro-caudal and dorso-ventral axes. The compiled MiSTR data provides novel insights into the earliest regionalised gene expression dynamics in the time period before establishment of neuralisation as well as the temporal expression patterns and external cues required for the formation of the human ventral forebrain and midbrain-hindbrain boundary (MHB).

## Results

### Cellular transcriptomic mapping of rostro-caudal and dorso-ventral fate acquisition

To study transcriptomic diversity across developing neural tube regions, we performed single cell and single nucleus RNAseq on microfluidic MiSTR tissues differentiated from the H9 hESC cell line, representing three different axes of the developing human neural tube: Model 1 representing the dorsal rostro-caudal axis (**R/C dorsal**), Model 2 representing the ventral rostro-caudal axis (**R/C ventral**) and Model 3 representing the dorso-ventral axis of the forebrain (**D/V forebrain**) (**Figure 1A-C**). All models were neuralised through dual SMAD inhibition from day 0 of differentiation (Chambers et al. 2009). The rostro-caudal tissues were induced by a 9-day microfluidic gradient of CHIR (WNTa), and global ventralisation was induced in the R/C ventral model through addition of SHH and purmorphamine. The D/V forebrain model was achieved through a WNT inhibition (WNTi) gradient with XAV939 from day 0-9, supplemented by a ventralising gradient comprised of SHH and purmorphamine from day 3-14 (**Figure 1B**). We developed a method to maintain the coherent MiSTR tissues in long-term cultures by covering the tissues with a layer of Matrigel on day 14 and then transferring the tissues from the device and into an orbital shaker. MiSTR tissues from all three models were collected for sc/snRNAseq at various timepoints from day 0 to day 62 (**Figure 1B**). In most cases, the tissues were sub-dissected into 5 regions across the gradient axis (A-E), and CITE-seq based hashtagging was applied to track the location of sequenced cells within the MiSTR tissue (Stoeckius et al. 2017) (**Figure 1C**). We sequenced a total of 277,513 cells/nuclei from 54 tissues covering 9 different timepoints with 2-3 biological replicates per timepoint, yielding a total of 206K cells after filtering and quality control (see **Figure S1A** for information on 10X experiments). From this, we obtained a total of 215 annotated clusters from analysing each timepoint between day 5 - 62 and each MiSTR model individually, and all of these clusters can be explored from https://cells.ucsc.edu/?ds=neural-tube-organoids on UCSC cell browser (Speir et al. 2021). To simplify the visualization of the dataset, we integrated the data from all three MiSTR models at all time points, thereby collapsing the dataset into 28 clusters encapsulating the overall rostro-caudal and dorso-ventral regionalization of the tissues (**Figure 1D-F** and **S1B-D**, cluster markers provided in **Supplementary Table S1)**. The dataset could be classified into three distinct phases of neural tube development: an early pre-neuralisation epiblast phase, characterized by markers such as *POU5F1, SOX2, HESX1*, and *ZIC3* **(Figure 1G)**, a regional neural progenitor phase, defined by markers such as *OTX2, SOX1, SOX2, PAX6, NKX2*.*1, HOXB1 and FGF17* **(Figure 1H)**, and a postmitotic neuronal phase, indicated by markers such as *STMN2, LHX6, ASCL1* and *PHOX2B* **(Figure 1I)**. Regional arrangement along the anterior to posterior axis was evident from markers such as *FOXG1, OTX2, NKX2-1, FGF17* and *HOXB1* (**Figure 1H**). Additionally, we verified by immunocytochemistry that the tissues gradually shifted from SOX2+ progenitors at d21 to MAP2+ neurons at d35 and that rostro-caudal regionality was maintained in tissues which had been cultured for 35 day, i.e. 26 days in the absence of gradient (**Figure 1J**). To explore neuronal diversity, we performed subclustering of the *STMN2* expressing cells and identified 24 distinct neuronal clusters displaying identities of various neurotransmitter phenotypes and regional identities, including cortical excitatory neurons, cortical interneurons, midbrain neurons, hindbrain neurons, hypothalamic neurons, retinal neurons and several others (**Figure 1K,L and Figure S2**).

**Figure 1.**
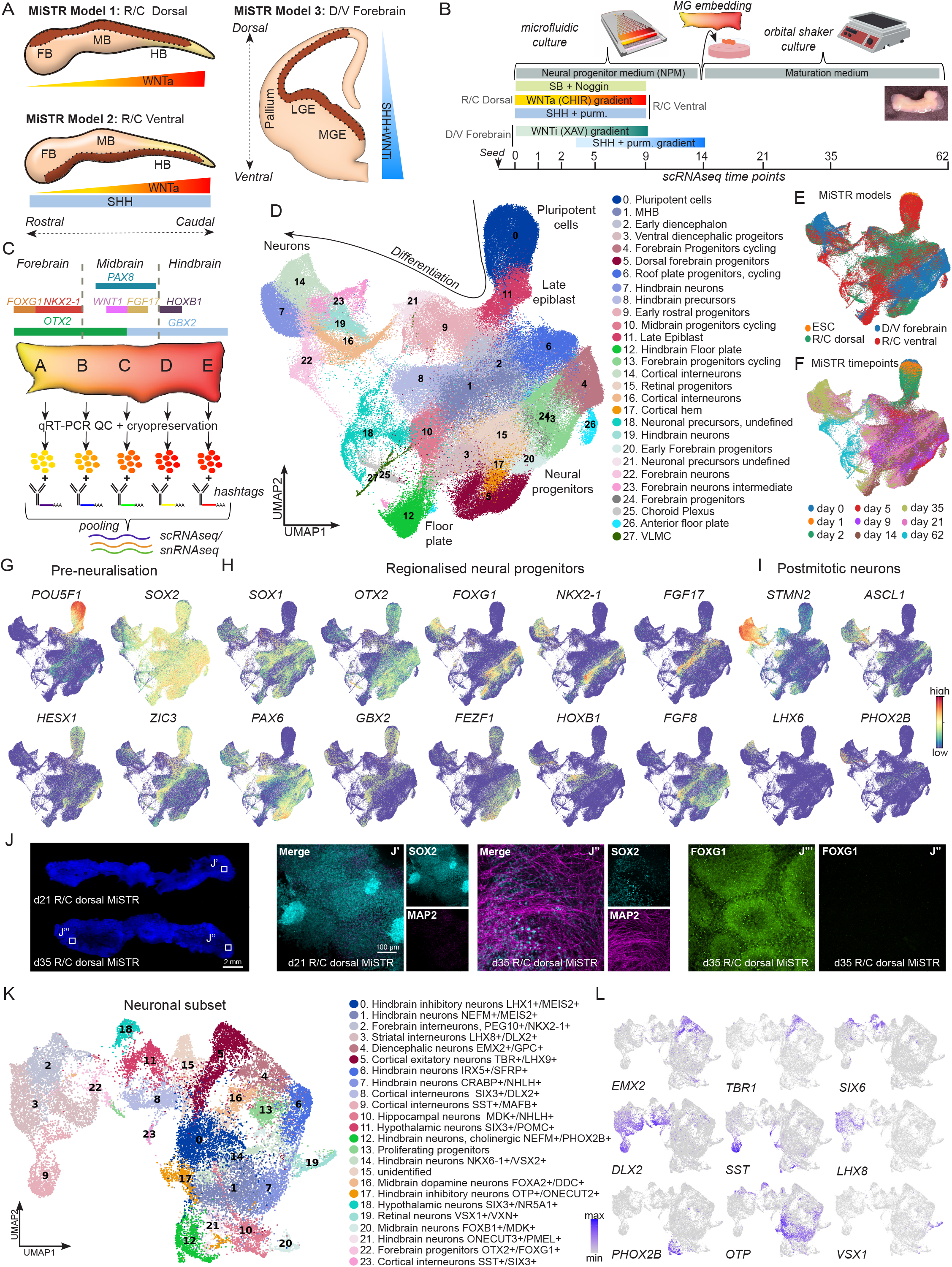
Integrated transcriptomic profiling of rostro-caudal and dorso-ventral MiSTR models. **A)** Schematic representation of the three different axes of the neural tube (in dark brown) that are modeled using the MiSTR system: The rostro-caudal axis was modeled through a WNT activation (WNTa) gradient (R/C dorsal), while adding global SHH to the WNTa gradient resulted in the generation of R/C ventral MiSTR tissue. Forebrain patterning was modelled through a SHH and WNT inhibition (WNTi) gradient (D/V forebrain). **B)** Overview of the protocol used to generate MiSTR tissues. Cells were differentiated in the microfluidic device for 14 days, and exposed to morphogen gradients for 9 days (R/C models) or 14 days (D/V forebrain model). For terminal maturation MiSTR tissues were embedded in Matrigel and cultured as organoids on an orbital shaker until day 62 (see inserted image of a d62 MiSTR tissue). The timeline indicates the timepoints when cells were harvested for sc/snRNAseq. **C)** Schematic of the gene expression pattern on a d14 rostro-caudal tissue, along with the strategy for sc/snRNAseq, where the MiSTR tissue was divided into five equal parts along the R/C or D/V axis (A-E regions). For each region at each timepoint, cells from 2-3 tissues were pooled in equal proportion and tagged with a unique hashtag antibody. The hashtagged cells from regions A-E were then pooled and analysed by 10X. **D)** UMAP embedding and annotated clusters of sc/snRNAseq data from all the sequenced MiSTR tissues; a dataset consisting of 9 timepoints (d0-62), 54 individual tissues (n = 2,3) and 206,246 cells. **E**,**F)** Equivalent UMAPs showing the three MiSTR models (E) and MiSTR tissue sequencing timepoints (F). **G**,**H**,**I)** UMAP plot of integrated MiSTR dataset showing the three development phases of MiSTR tissue, an early pre-neuralisation phase (marked by *POU5F1*(*OCT4*), *SOX2, HESX1* and *ZIC3*), regional neural progenitor phase (*OTX2, SOX1, PAX6, NKX2*.*1, HOXB1 and FGF17*) and postmitotic neuronal phase (*STMN2, LHX6, ASCL1* and *PHOX2B*). **J)** Whole mount confocal images from R/C Dorsal MiSTR tissues at d21 and d35 (left panel). SOX2 and MAP2 immunocytochemistry (middle panel) of d21 (**J’**) and d35 (**J’’**) R/C dorsal MiSTR tissues, showing matured MAP2^+^ neurons at d35, and FOXG1 immunostaining (right panel), present in the rostral end (**J’’’)** of d35 R/C dorsal MiSTR tissue, but not in the caudal end (**J”**), thus showing that rostro-caudal regionality was maintained in tissues which had been cultured for 35 days. **K-L)** UMAP plot of only neuronal (high *STMN2* expressing) cells from the integrated MiSTR dataset, depicting different neuronal clusters along with gene expression plot for forebrain neurons, interneurons, retina and hindbrain (28,402 cells).

### MiSTR tissues map to specific axes of the early human neural tube

To assess the reliability of the MiSTR models in representing early human neural tube development, we mapped our data from the three MiSTR models to a comprehensive scRNAseq dataset of the first trimester of human fetal brain development(Braun et al. 2023). Deep learning label transfer facilitated this comparison using scvi-tools (Xu et al. 2021, Gayoso et al. 2022). The label transfer showed that gradient-patterned cells from the R/C dorsal and R/C ventral MiSTR tissues exhibited similarity to fetal cells of all forebrain, midbrain, and hindbrain identities of developing human brain between PCW 5.5 to 9. In contrast, cells from the D/V Forebrain model contained only cells with similarity to fetal forebrain (**Figure 2A, B and S3A-C**). To visualise spatially the regionalisation covered by the three models, we further mapped MiSTR cells from all three models at day 14 and 21 to the spatial transcriptomic dataset of human fetal brain at 5 PCW by employing Bonefight and Tangram (Biancalani et al. 2021, La Manno et al. 2021, Braun et al. 2023). This mapping demonstrated alignment of R/C dorsal MiSTR tissues to the dorsal axis of the neural tube, R/C ventral MiSTR tissues to the ventral axis of the neural tube, and D/V forebrain MiSTR tissues exclusively to the forebrain region of the neural tube (**Figure 2C**). We concluded from these findings that the three MiSTR models, based on patterning gradients targeting only two morphogenic pathways, WNT and SHH, effectively recapitulated early human neural tube patterning from forebrain to hindbrain along both the rostro-caudal and dorso-ventral axes.

**Figure 2.**
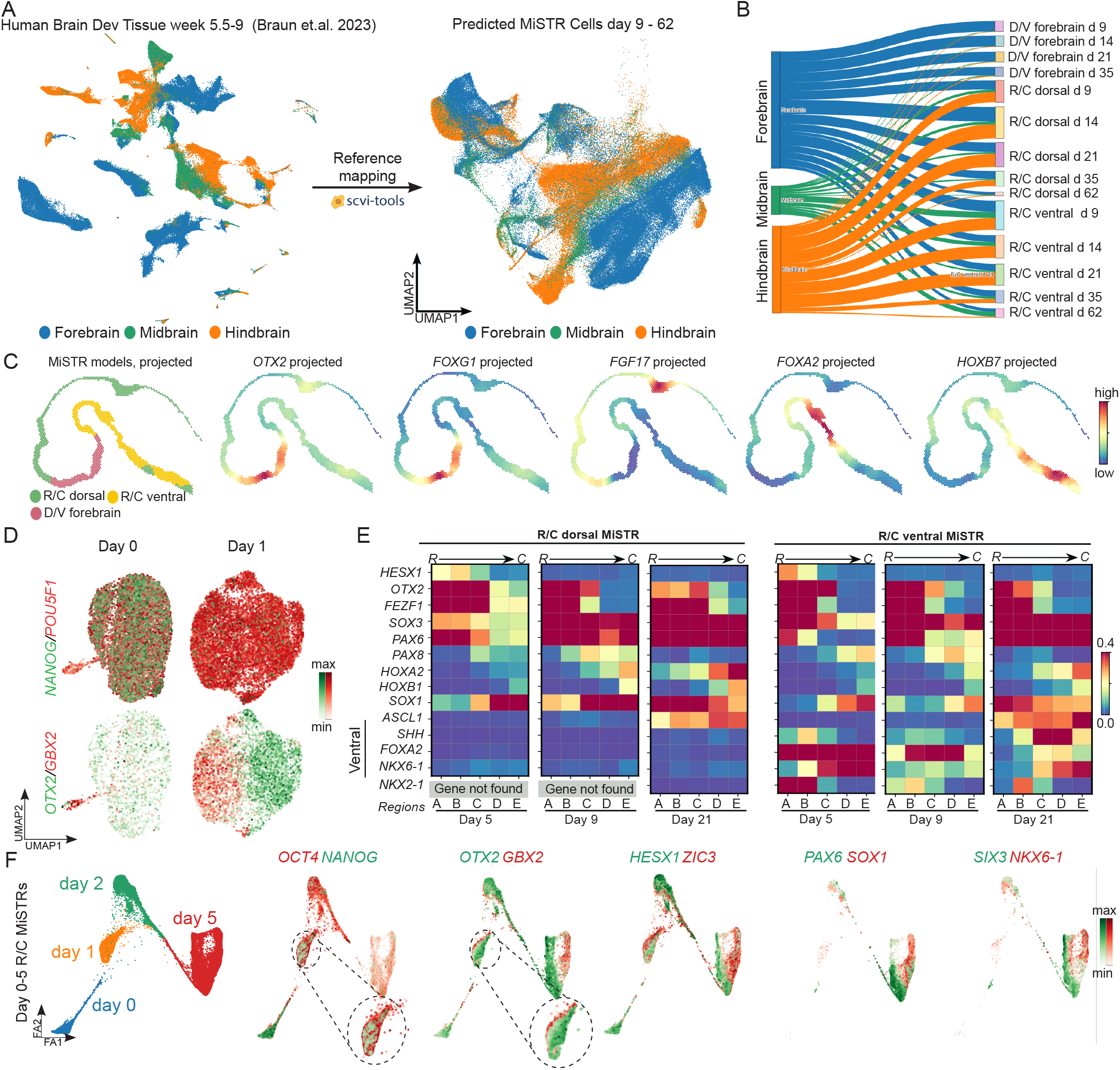
Mapping regional and temporal expression dynamics in MiSTR models. **A)** Developing human fetal brain atlas from week 5 -9 (Braun et al. 2023) plotted with original publication embedding with annotations compiled to Forebrain, Midbrain and Hindbrain (left side) and UMAP embedding of the fetal annotations mapped onto the integrated MiSTR dataset (d9-62) (right side). **B)** A Sankey diagram depicting the proportion of cells from each MiSTR timepoint and model mapped with Forebrain, Midbrain and Hindbrain cells of the human brain atlas. **C)** Spatial mapping of integrated d14 and d21 MiSTR tissues from all there MiSTR models projected onto the PCW 5 human fetal neural tube (Braun et al. 2023). The projected expression of regional markers OTX2, FOXG1, FGF17, FOXA2 and HOXB7 in the mapped MiSTR voxels validated the mapping against the original transcriptomic measurements in the embryo. **D)** UMAP plots showing the expression of pluripotency markers *POU5F1* (OCT4) and *NANOG*, as well as rostro-caudal (R/C) segregation markers *OTX2* and *GBX2* in d0 human embryonic stem cells (hESCs) and d1 R/C ventral MiSTR tissue. **E)** Matrix plots showing temporal dynamics of region specific markers in the five regions (A-E) of R/C dorsal and R/C ventral MiSTR tissues at different timepoints (day 5, 9 and 21). The establishment of rostro-caudal pattering is visualized by expression of *FOXG1, OTX2, PAX8* and *HOX* genes, while emergence of ventral fates in R/C ventral MiSTRs are shown by expression of *SHH, NKX2-1, FOXA2* and *NKX6-1*. **F)** Force atlas2 (FA2) embeddings of d0-5 R/C MiSTRs showing that rostro-caudal patterning is established already at d1-2, marked by *OTX2* and *GBX2* expression, followed by establishment of neural identity at d5 as marked by the expression of *PAX6* and *SIX3* in the rostral domain and *SOX1* and *NKX6-1* in the caudal domain (only cells expressing gene are shown).

We next compared the temporal dynamics of region-specific markers in scRNAseq data from the five hashtagged regions (A-E) of R/C dorsal, R/C ventral and D/V forebrain MiSTR tissues (see experimental outline in **Figure 1A-C**). We confirmed previous observations (Rifes et al. 2020) that rostro-caudal patterning was established very early in the R/C MiSTRs, as evident by opposing expression patterns of *OTX2* versus *GBX2* by 24 hours of differentiation (**Figure 2D**), and *OTX2* versus *HOXA2/HOXB1* by d5-9 of differentiation in the rostral versus caudal regions (**Figure 2E**). As expected, the R/C ventral MiSTR model further exhibited enrichment of ventral markers (*FOXA2, SHH* and *NKX6-1*), while these markers were absent in the R/C dorsal MiSTR model (**Figure 2E**). Analysis of cells from the earliest time points (day 0-5) showed that while the rostro-caudal specification of OTX2+ and GBX2+ fates occurred already at 24 hours of differentiation, key transcriptional regulators of neural fate (i.e. SOX1 and PAX6) did not turn on before day 5 of differentiation (**Figure 2F**). More surprisingly, this analysis also showed that neuralisation progressed through different transcriptional programs in the rostral versus caudal domains, with *PAX6* being dominantly expressed in the *OTX2*^+^ population and *SOX1* being dominantly expressed in the *GBX2*^+^ population at day 5 of differentiation (**Figure 2F**).

### Neuralisation of rostral (OTX2^+^) and caudal (GBX2^+^) fates occurs through divergent transcriptional programs

Next, we explored the dynamics of rostro-caudal fate establishment more closely through integration and pseudotime analysis of isolated *OTX2*^+^ and *GBX2*^+^ populations from day 0-9 of differentiation. This analysis confirmed clear differences in the transcriptional programs initiated in the rostral versus caudal populations during the earliest stages of differentiation. The *OTX2*^+^ population showed an early expression wave of *HESX1, CYP26A1* and *SHISA2*, whereas the *GBX2*^+^ population in contrast showed a strong induction of *ZIC3, SP5* and *FST* during the first days of patterning (**Figure 3A,B**). According to the pseudotime analysis, these early rostro-caudal patterning events occurred during a time window at which the cells still expressed *OCT4* (*POU5F1*), and before the onset of expression of neuroepithelial markers *PAX6* and *SOX1* (**Figure 3B**). The analysis further confirmed that while *PAX6* emerged as the first neuroepithelial marker appearing in the rostral (*OTX2*^+^) domain, *SOX1* was simultaneously activated exclusively in the caudal domain (*GBX2*^+^) (**Figure 3B**). To complement these findings from the MiSTR with an alternative model, we conducted 2D monolayer differentiations with an alternative cell line (KOLF2.1J hiPSC line). The cells in the monolayer culture were differentiated towards either forebrain (0 μM CHIR) or hindbrain (2 μM CHIR) fates (**Figure 3C**). Through temporal qRT-PCR analysis from d0-9, we confirmed a strong activation of *OTX2, HESX1, SHISA2, FEZF1* and *LHX5* in the forebrain differentiations during the early pre-neuralisation phase from d0-3. In contrast, *GBX2, FST, HOXA1* and *CRABP2* were strongly upregulated in the hindbrain differentiations during this phase. The pre-neuralisation phase was followed by activation of *PAX6* and *DLK1* in the forebrain-patterned cells (Figure 3D) and *SOX1* and *HOXA2* in the hindbrain-patterned cells around d4 (**Figure 3E**), thereby confirming our findings from the MiSTR. To assess the in vivo relevance of our findings from the preneuralisation stage, we mapped early time points of MiSTR tissue development (days 0, 1, 2, and 5) to an in vivo dataset. Since high-resolution temporal datasets are not available from the human embryo at these early stages, we employed a temporally dense dataset from the gastrulating mouse embryo E7.0-E8.5 (Pijuan-Sala et al. 2019). As expected, the majority of cells at d0 and d1 were predicted to be epiblast cells, whereas increasing prediction of rostral and caudal mouse neurectoderm emerged at d2 (**Figure S4A-C**) despite the strong retained expression of *OCT4* and lack of *PAX6* and *SOX1* in the human cells at this time point (**Figure 3C-E**). Furthermore, gene expression visualisation in the mouse populations of Epiblast, Caudal epiblast, Rostral Neurectoderm, Caudal Neurectoderm from E7.0 to E8.0 showed similar expression pattern of pre-neuralisation markers (**Figure S4D**). Taken together, these findings confirm that the pre-neuralisation phase in the mouse embryo is characterised by expression of a similar set of rostral and caudal fate genes, suggesting that neural tube regionalisation in both human and mouse occurs significantly earlier than the onset of neuralising transcription factors.

**Figure 3.**
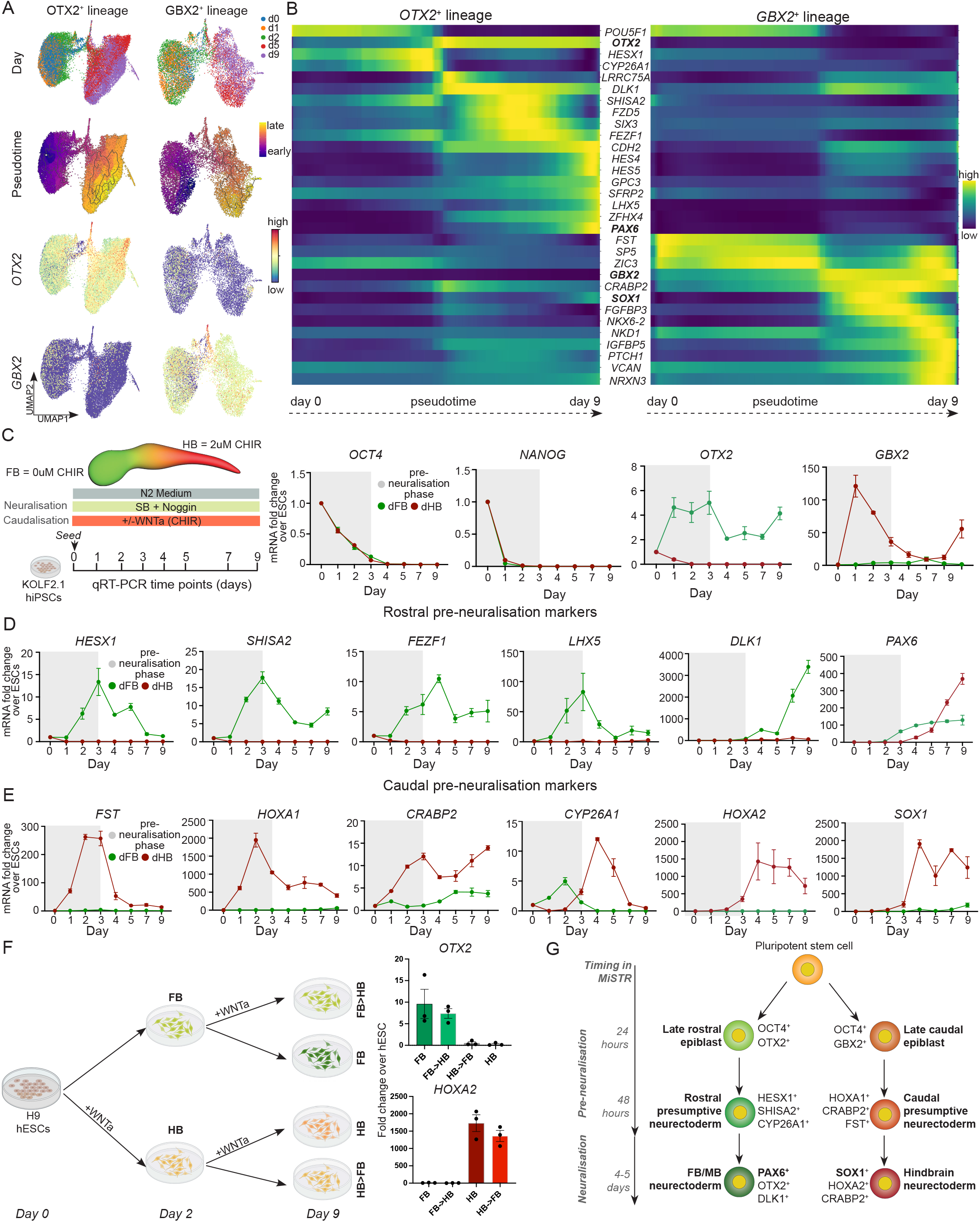
Decoding divergent transcriptional programs for rostral (OTX2^+^) and caudal (GBX2^+^) fate specification of early neural plate development. **A)** UMAP plots of OTX2+ and GBX2+ cells, along with pluripotent cells (d0), showing the progression of pseudotime from d0 to 9. **B)** Heatmap of OTX2+ and GBX2+ cells from d0 to 9 along the pseudotime axis showing the differential expression pattern of key genes involved in patterning of rostral vs caudal neural population. **C)** Schematic of forebrain (FB) and hindbrain (HB) patterning of hiPSCs (KOLF 2.1 line) along with qRT-PCR plots of *NANOG, OCT4, OTX2 and GBX2* (n = 3) (pre-neuralisation in gray refers to state of cells). **D-E)** qRT-PCR based fold change plots of rostral and caudal pre-neuralisation markers in dorsal forebrain (dFB) and dorsal hindbrain (dHB) lineages and their differential expression throughout the early neural patterning process (n = 3). **F)** Schematic of the 2D cell culture assay designed to investigate the impact of CHIR dosage manipulation on hESC patterning towards dFB and dHB lineages followed by qRT-PCR analysis of *OTX2* and *HOXA2* to confirm rostral and caudal identities (n=3). **G)** Schematic based on early MiSTR patterning scRNAseq data proposing two divergent transcription factor programs resulting in rostral and caudal patterning of the early neural plate. Pluripotent stem cells initially differentiate into two distinct types of regionally patterned epiblasts: late rostral epiblast (expressing *OCT4* and *OTX2*) and late caudal epiblast (expressing *OCT4* and *GBX2*). This is followed by a rostro-caudal patterning event, stemming in differentiation of the late rostral epiblast into rostral presumptive neuroectoderm (*HESX1, SHISA2* and *CYP26A1*) eventually leading to the formation of Forebrain / Midbrain neurectoderm (*PAX6,OTX2* and *DLK1*). Similarly, late caudal epiblasts differentiate into caudal presumptive neurectoderm (*HOXA1,CRABP2* and *FST*) eventually propagating towards hindbrain neurectoderm (*SOX1,HOXA2* and *CRABP2*).

To test if the OTX2^+^ rostral and GBX2^+^ caudal fates were fate-determined already during the pre-neuralisation phase, we conducted a cross-over experiment in which we differentiated monolayer cultures of H9 hESCs and exposed them to either forebrain (0 μM CHIR) or hindbrain (2 μM CHIR) patterning conditions for 2 days, after which the cells were either continued in the same medium or switched to the reverse CHIR concentration from d2 to d9 of differentiation.This experiment revealed that two days of initial WNT activation by CHIR was equally efficient at inducing hindbrain fates (loss of *OTX2* and induction of *HOXA2*) as 9 sequential days of CHIR stimulation (**Figure 3F**). Similarly, the reverse experiment showed that cells which were exposed to two days of forebrain patterning could no longer be induced to hindbrain fates when exposed to 2 μM CHIR from day 2-9 (**Figure 3F**). This data shows that rostral and caudal fate specification is fully established during the first 2 days of WNTa exposure, before the onset of PAX6 and SOX1 expression. Based on this data, we propose a novel neural bifurcation model in which epiblast cells first bifurcate into *OCT4*^*+*^/*OTX2*^*+*^ late rostral epiblast and *OCT4*^*+*^/*GBX2*^*+*^ late caudal epiblast **(Figure 3G, top)**. This is in the rostral OTX2^+^ epiblast followed by expression of presumptive rostral neuroectoderm markers *HESX1, LHX5, SHISA2*, and *CYP26A1*, which may serve to inhibit caudalisation through their known functions in blocking WNT, RA and FGF signalling, respectively (Nagano et al. 2006, Peng and Westerfield 2006, Thatcher and Isoherranen 2009, Andoniadou et al. 2011) (**Figure 3G, middle**). The pre-neuralisation phase of the presumptive caudal neurectoderm is in turn marked by the expression of CRABP2 which enables responsiveness to retinoic acid (RA) (Dong et al. 1999) and FST which inhibits BMP signalling (Amthor et al. 2002). The presumptive neuroectoderm cells subsequently undergo proper neuralisation with PAX6 as the pioneering neuralising factor in the rostral cells and SOX1 as the neuralising factor in the caudal cells **(Figure 3G, bottom)**.

### Forebrain ventral transcriptional programs occur only after neuralisation

We next turned towards performing a similar analysis of the temporal dynamics of ventralisation in the D/V forebrain MiSTR (**Figure 4A**). While the R/C MiSTRs showed clear rostro-caudal regionalization already by day 1 (**Figure 2D**), the early D/V forebrain MiSTR showed no signs of dorso-ventral regionalization even at day 5 of differentiation, despite 2 days of prior exposure to a SHH/purmorphamine gradient (**Figure 4B**). Analysis of hashtagged subdissected regions in the tissues showed global co-expression of dorsal and ventral markers (i.e. *PAX6, SIX3, SIX6* and *RAX*) across the entire tissue at day 5-9 (**Figure 4B**). In contrast, integration of all timepoints from the D/V forebrain model confirmed that dorso-ventral fates were established later in the tissue, giving rise to both pallial and subpallial progenitors (**Figure 4C-E**). Unbiased comparison of the global transcriptome of d5 subsetted cells from the high SHH regions (D/E) versus low SHH regions (A/B) confirmed no detectable transcriptomic differences in these cell populations despite 2 days of exposure to a SHH/purmorphamine gradient in contrast to the strong D/V difference apparent at this stage following SHH in the R/C patterning conditions (**Figure 4F**). Even at day 9, apart from a small distinct cluster of diencephalic cells (*FOXD1*^+^), the ventral transcription factor *NKX2-1* was not yet activated in the telencephalic population, even though cells had at this point been exposed to SHH-purmorphamine gradient for 6 days (**Figure 4G,H**). The D/V Forebrain model included a 3-day rostralisation step with a WNTi gradient (XAV939) to inhibit formation of ventral diencephalic fates (**Figure 1A**), and the highly delayed ventralisation in this model was in stark contrast to the R/C ventral MiSTR which had not been exposed to WNTi treatment and which displayed strong and widespread expression of both *NKX2*.*1* and *FOXA2* already by day 5 (**Figure 2E**). This indicated that temporal permissiveness to ventralising cues differed between the most rostral telencephalic domain exposed to WNTi (D/V forebrain MiSTR) and the more caudal diencephalic/midbrain domains of the R/C ventral MiSTR.

**Figure 4.**
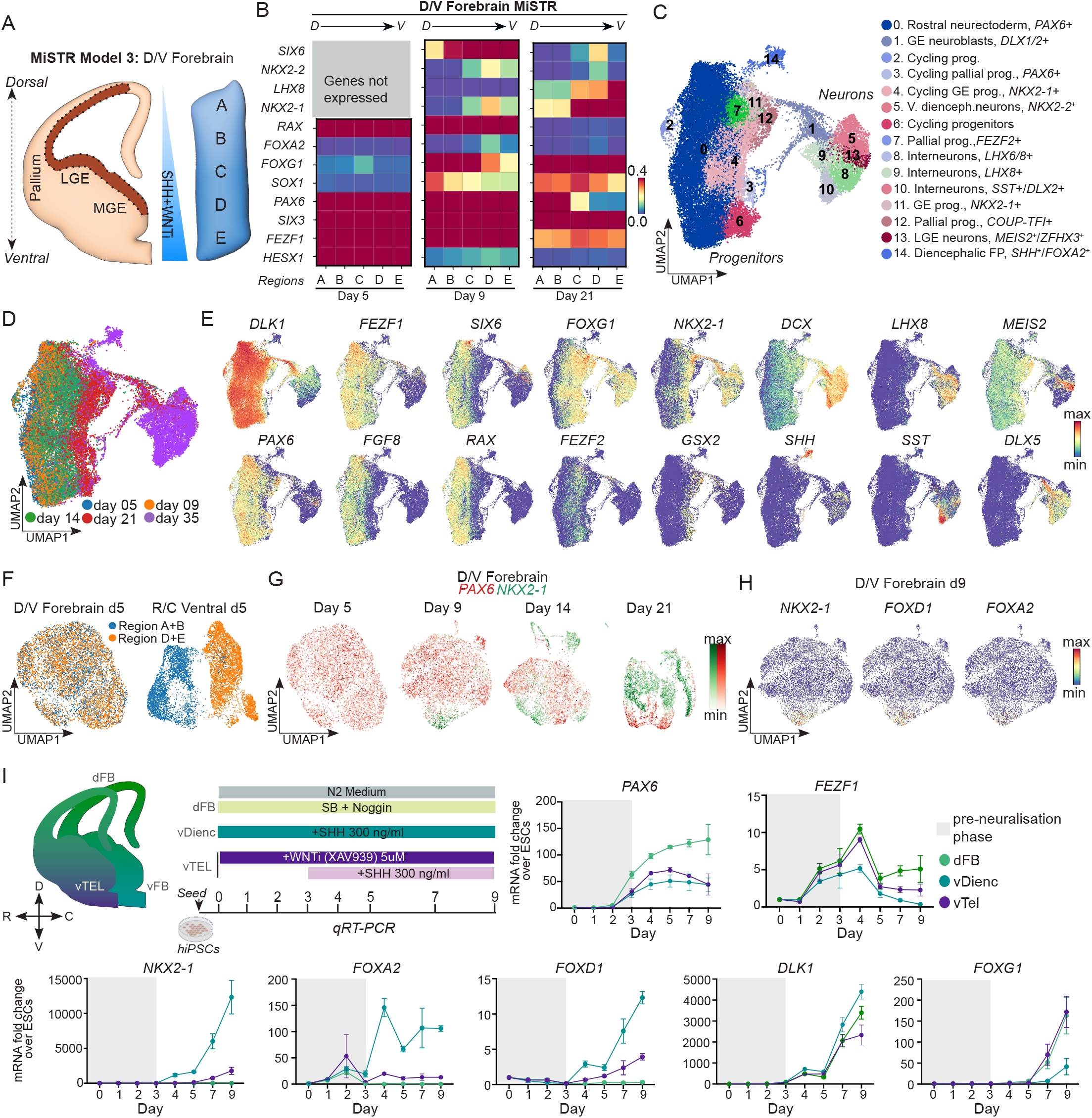
Development dynamics of early telencephalic patterning using D/V forebrain MiSTR tissue. **A)** Schematic depicting the areas of the anterior telencephalon modeled by D/V forebrain MiSTR tissue (in dark brown), where the A region represents the most dorsal side and the E region represents the most ventral side. **B)** Matrix plots from scRNAseq data showing temporal dynamics of region-specific markers in D/V forebrain MiSTR tissues at day 5, 9 and 21. By day 21, all regions express *FOXG1* (a pan-telencephalic marker). PAX6 marks the dorsal territory in regions A and B, while *NKX2-1* marks the ventral territory in regions D and E. **C**) UMAP embedding of integrated timepoints (d5, 9, 14, 21, 35) from D/V forebrain MiSTR with the cluster annotations. **D-E**) Equivalent UMAP showing the arrangement of the cells from these time points, organizing themselves from the left side (d5) to the right side (d35). This marks the progression of late rostral epiblast cells to various anterior telencephalic subtypes, supported by gene plots. *FGF8* marks the earliest phase of early forebrain patterning, later replaced by *FOXG1*, accompanied by various telencephalic subtype markers such as *SST, DLX5, LHX8*, and *MEIS2*. **F)** UMAP plot of individual timepoints of D/V forebrain MiSTR tissue depicting the expression domains of *PAX6* and *NKX2-1*. As early as day 5, the entire tissue was marked by *PAX6* showing no signs of Dorso-Ventral regionalisation despite 5 days of WNTi and 2 days of SHH exposure, subsequently the expression of *NKX2-1* was initiated at day 9, eventually establishing distinct dorsal and ventral territories by day 21. **G)** UMAP plots of d5 R/C Ventral MiSTR and D/V forebrain MiSTR coloured by A+B, and by D+E regions, showing that WNT activation in the R/C Ventral MiSTR elicits a rostro-caudal regionalisation already by d5, while WNT inhibition in the D/V forebrain MiSTR does not result in a dorso-ventral regionalisation. **H)** UMAP plots of d9 D/V forebrain MiSTR showing a small isolated cluster marked by the diencephalic marker *FOXD1* also expressing *NKX2-1*. **I)** Schematic strategy of patterning hiPSCs (KOLF2.1J) in three condtions namely dorsal Forebrain, Ventral diencephalon and ventral telencephalon along with qRT-PCR based fold change expression of key telencephalic *(FOXG1)* and diencephalic (*FOXD1*) markers (n = 3).

To test this, we conducted 2D monolayer differentiations with KOLF2.1J hiPSCs patterned towards dorsal forebrain (dFB) (SB/Noggin), ventral telencephalic (vTel) fates (SB/Noggin + XAV393 from day 0 + SHH from day 3) and ventral diencephalic fates (vDienc) (SB/noggin + SHH from day 0). We found the telencephalic marker *FOXG1* to be enriched in the vTel cultures while the diencephalic marker *FOXD1* was upregulated in the vDienc cultures, confirming the regional patterning of the cultures (**Figure 4I**). Consistent with findings from the MiSTR, we observed early upregulation of ventral markers *NKX2-1* and *FOXA2* by day 4 in the vDienc cultures, while in the vTel condition, upregulation of ventral markers was delayed until day 9, and was much less prominent (**Figure 4I**). The onset of NKX2.1 expression in the vTel condition coincided with activation of the telencephalic marker FOXG1.

In summary, data from both the MiSTR and from monolayer differentiations indicated that neural ventralisation occured significantly later than rostro-caudal specification and that the ventralizing program was even further delayed in the most rostral ventral telencephalic lineage compared to the ventral diencephalic linage.

### Early cell fate bifurcation in human ventral forebrain development

We next investigated the dynamics of the more rapid ventralising response in the diencephalic regions of the R/C ventral MiSTR. We observed from the scRNAseq data and immunofluorescence images, that *OTX2, NKX2-1, and FOXA2* were broadly co-expressed in the day 5 R/C ventral MiSTR (**Figure 5A-C)**. In contrast, d9 and d14 tissues showed loss of the co-expressing population and emergence of mutually exclusive populations of *NKX2-1*^+^ and *FOXA2*^+^ cells (**Figure 5C**). To understand the lineage bifurcation events underlying the segregation of the *FOXA2*^*+*^*/NKX2-1*^*+*^ population into *NKX2-1*^*+*^ and *FOXA2*^*+*^ single-positive populations, we subsetted all cells from day 5 to day 14 expressing either *NKX2-1* or *FOXA2*, while filtering away all *OTX2*-negative hindbrain cells. In this new subsetted dataset (**Figure 5D)**, we searched for active gene regulatory networks (GRNs) using SCENIC (Van de Sande et al. 2020) in each of the three main subtypes: *NKX2-1*^*+*^ and *FOXA2*^*+*^ single positive cells and *NKX2-1/FOXA2* co-expressing cells (**Figure 5E)**. We next performed *in silico* gene perturbation of the top identified regulons in the 3 populations using CellOracle (Kamimoto et al. 2023). The knock-out (KO) simulations targeting *SIX3, RAX* and *TCF7L2* showed suppression of the *NKX2-1*^*+*^ forebrain domain, and a transition towards the domain expressing *FOXA2* (**Figure 5F)**. In contrast, the *IRX1* KO simulations showed severe disruptions in the differentiation of cells at day 9 and 14, with a regression back towards the precursor *NKX2-1*^+^/*FOXA2*^+^ cell state, suggesting a role for IRX1 in lineage birfurcation of the early co-expressing population towards diencephalic and midbrain identities (**Figure 5F**). IRX1 and its isoform IRX1b have been reported in diencephalic development and establishemnt of Zona Limitans Intrathalamica (Scholpp et al. 2007, Li et al. 2010). In summary, we propose from our observations a model where the early ventralisation in the telencephalon is blocked by pre-exposure to WNTi, whereas early ventralisation of the diencephalon is permissive and involves the initial formation of co-expressing NKX2-1^+^/FOXA2^+^ cells which subsequently birfucate into single-positive populations, potentially through the actions of IRX1 (**Figure 5G**).

**Figure 5.**
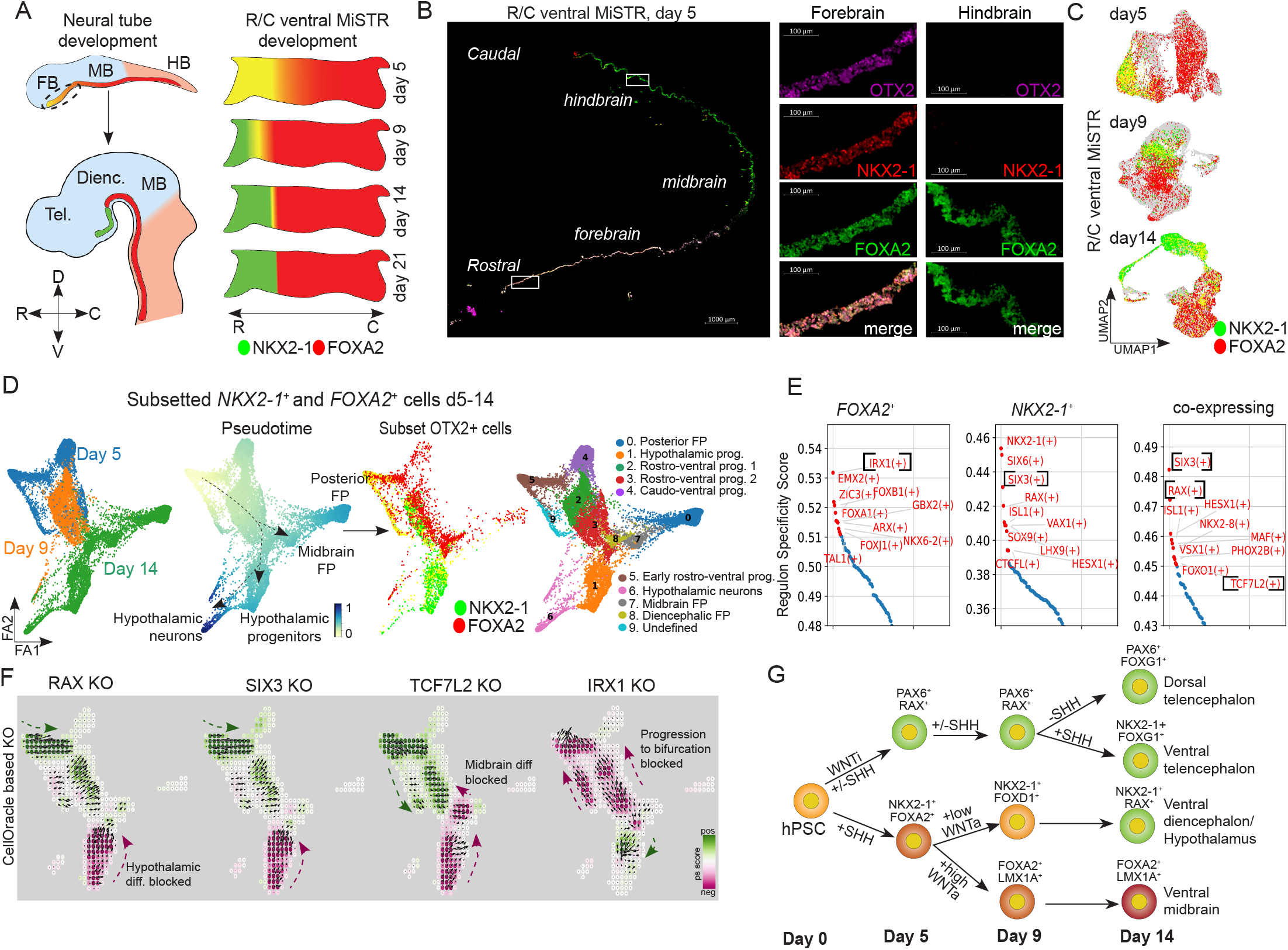
Development dynamics of early telencephalic/diencephalic patterning using MiSTR tissue. **A)** A schematic showing the expression dynamics of *FOXA2* and *NKX2-1* in the developing neural tube and R/C ventral MiSTR tissue over time. **B)** Immunofluorescence microscopy image of sectioned d5 R/C ventral MiSTR tissue showing the co-expression of *NKX2-1* and *FOXA2* in the rostral side of tissue marked by expression of *OTX2*. Scale Bars 1000 μm and 100 μm. **C)** UMAP embedding of *NKX2-1*^+^, *FOXA2*^+^ and double positive cells from d5 to 14 in R/C ventral MiSTR, depicting the gradual decrease in double positive cells over time. **D)** Cells expressing *NKX2-1* and/or *FOXA2* from d5, 9 and 14 in R/C ventral MiSTR were subsetted and plotted on FA2 embedding along with their projected pseudotime and cluster annotations. This dataset was further subsetted to only contain *OTX2* positive cells to remove hindbrain populations from the dataset. **E)** Top SCENIC regulons were identified specifically in the *FOXA2*^+^ population, the *NKX2-1*^+^ population and the co-expressing population from the subset dataset. **F)** Celloracle knockout simulations of the selected transcription factors *RAX, TCF7L2, SIX3, IRX1* and *POU3F1* depicting positive and negative differentiation scores along with the transition vectors. **G)** Schematic digram of progression of MiSTR developmet in dorso-ventral telencephalic lineage along with ventral diencephalic/hypothalamus and midbrain lineage.

### Gradients of only SHH and WNT can generate secondary signalling centers

We next investigated whether the initial 9 and 14 days of morphogen gradient patterning were sufficient not only to regionalize cell types along the rostro-caudal and dorso-ventral axes but also to induce secondary signalling centers that secrete local morphogens. Heatmap visualization of developmental morphogens and their respective receptors extracted from scRNAseq clusters compiled from all three models (**Figure 1D**) showed highly specific regional expression patterns, including the co-expression of *FGF8, 17*, and *19* in the midbrain and MHB clusters, as well as localized expression patterns of *WNT5A, BMP1, SHH, SLIT1/2/3*, and *PDGFC* in the floor plate clusters (**Figure 6A and S5B**). In addition, we observed distinct expression of *WNT7A, WNT7B, RSPO1, RSPO2*, and *RSPO3* in the Forebrain cluster **(Figure 6A and S6B)**. This demonstrated that, despite limiting exogenous patterning to only two morphogen gradients, WNT and SHH, multiple secondary signalling centers with region-specific secreted molecules were efficiently established within the MiSTR tissues, including the MHB.

**Figure 6.**
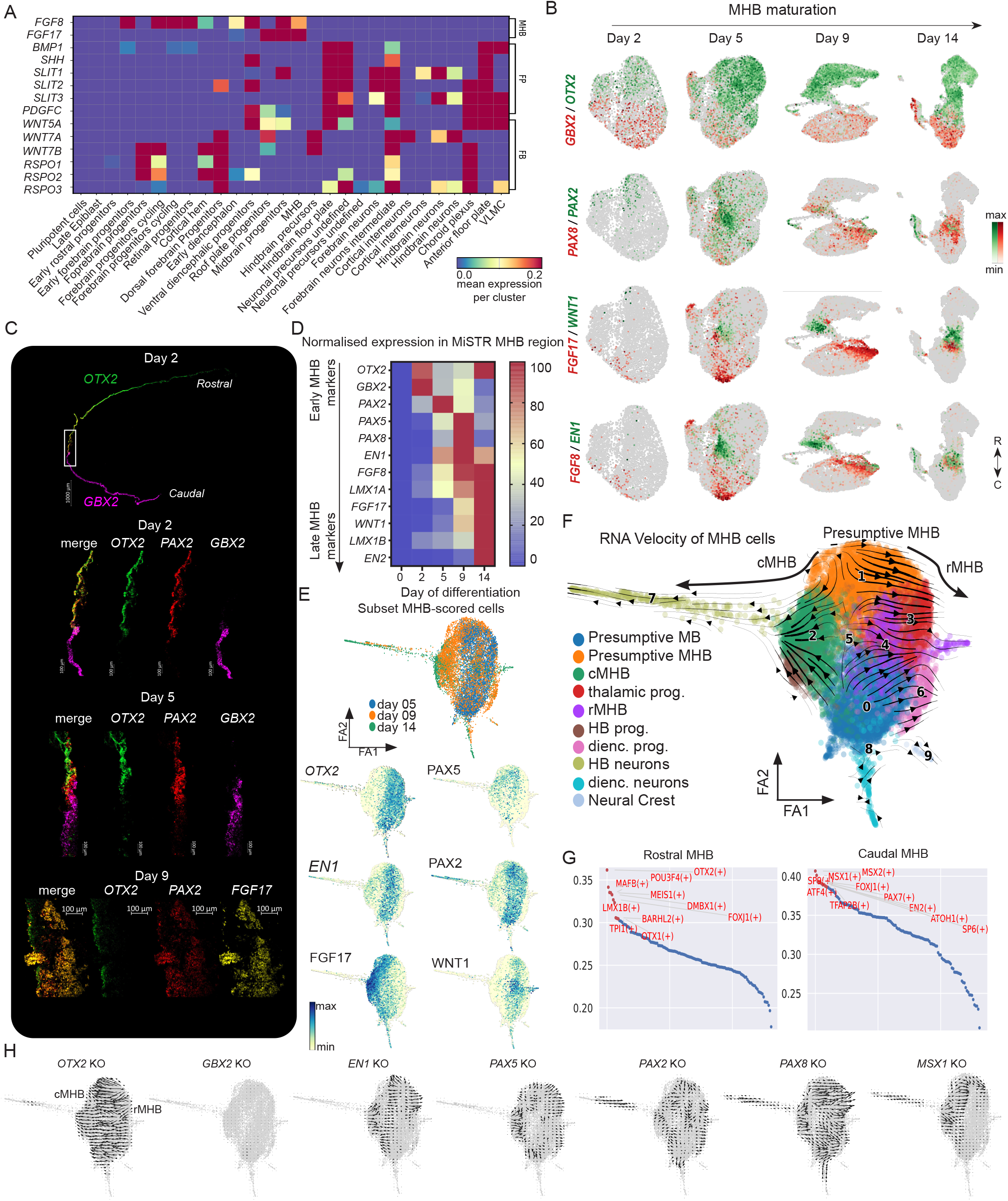
Expression dynamics of early and late MHB markers. **A)** Matrix plot of secreted growth factors across selected clusters of the integrated MiSTR dataset from all three models (d0-62), showing the emergence of region specific secondary signalling centers in the developing MiSTR tissues such as isthmic organizer, floor plate and forebrain. **B)** Gene expression plots showing the regional markers OTX2 (rostral) and GBX2 (caudal) together with MHB markers FGF17, WNT1, PAX2, PAX8, FGF8, and EN1 in developing R/C dorsal MiSTR tissues on day 2, 5, 9, and 14. **C)**RNAscope *in situ* hybridisation image of d2 R/C dorsal MiSTR showing *OTX2*^+^ and *GBX2*^+^ expression domains, the juxtapositioning of *OTX2* ad *GBX2* expression marks the location of MHB development. *In situ* hybridization imaging of *OTX2, PAX2* and *GBX2* in the presumptive MHB zone (region C) in d2 and 5 R/C dorsal MiSTR tissues showing *PAX2* expression spanning both the *OTX2*^+^ and *GBX2*^+^ side of the MHB. Equivalent RNAscope for *OTX2, PAX2* and *FGF17* in d9 R/C dorsal MiSTR tissues shows *PAX2* expression at d9 overlapping with both the *OTX2*-expressing and the *FGF17*-expressing domains. Scale Bars 1000 μm and 100 μm. **D)** Heatmap of qRT-PCR based normalized expression of MHB markers from MiSTR C-region in day 0, 2, 5, 9 and 14 tissue showing early and late markers of MHB formation in the MiSTR system (n = day 2: 5-10, day 5: 3, day 9: 5, day 14: 12-22). **E)** Forceatlas (FA2) embedding of MHB cells from day 5, 9 and 14 (see methods) showing the organisation of cells from d5 to 14 from center to outwards respectively along with key MHB markers. **F)** RNA velocity stream vectors obtained from the subset MHB dataset embedded on FA2 embedding layout depicting the directionality of cellular differentiation along with cluster annotations. **G)** Top regulons enriched by SCENIC analysis specific to each cluster in the MHB subset dataset. **H)** The transition vectors obtained from the CellOracle-based KO simulation plots indicating the shift in cell identity of in-silico knock-out candidate genes *OTX2, GBX2, EN1, PAX5 PAX2, PAX8 and MSX1*.

### Temporal dynamics of MHB establishment

We focused our analysis further on the MHB, an early transient secondary organiser which plays a crucial role in subregional patterning of the midbrain and hindbrain and which has been inherently difficult to study in humans. Temporal scRNAseq and in situ hybridization showed the early segregation of *OTX2*^+^ and *GBX2*^+^ domains at day 2 followed by expression of *PAX2, PAX8* and *EN1* at the OTX2/GBX2 intersect on day 5 and emergence of segregated rostral MHB (*OTX2*^*+*^/*WNT1*^*+*^) and caudal MHB (*GBX2*^*+*^*/FGF8*^*+*^*/FGF17*^*+*^) populations at day 9, with *EN1* and *PAX8* being expressed on both sides of the boundary (**Figure 6B-D**). The rostral and caudal MHB was maintained at day 14, although the boundary was at this point characterised mainly by expression of *WNT1* and *FGF17*, while *FGF8* was downregulated (**Figure 6B)**. Temporal mapping of gene expression patterns in the C-region of the R/C MiSTR (i.e. the region harbouring the presumptive MHB) confirmed *PAX2* as the earliest marker of the presumptive MHB at d5, followed by expression of *PAX5, PAX8, EN1* and *FGF8* at day 5-9 and subsequently by *WNT1, FGF17, EN1* and *EN2* marking the mature MHB structure at day 14 (**Figure 6D**). In order to investigate the active GRNs within the emerging MHB populations over time, we subsetted all cells with a positive cumulative MHB score from 5, 9 and 14 of R/C dorsal MiSTRs based on the expression of *PAX2, PAX8, EN1, FGF8, FGF17* and *WNT1* (**Figure 6E**, see also Methods). From this, we obtained 10 distinct clusters associated with different time points of MHB maturation (**Figure 6F**). RNA Velocity analysis showed two clearly opposing differentiation trajectories. The early *OTX2*^+^ population showed a trajectory towards the rostral MHB and subsequently towards midbrain and thalamic neurons. The early *GBX2*^+^ population showed a trajectory towards the caudal MHB and subsequently towards hindbrain neurons (**Figure 6F**). To assess the importance of key MHB gene regulatory networks, we performed SCENIC analysis of the rostral versus caudal MHB populations and proceeded to perform *in-silico* knockout simulation of selected transcription factors using CellOracle (Kamimoto et al. 2023). We found that OTX2 was the top transcription factor regulon identified in the rMHB cluster (**Figure 6G**), and indeed, simulated knockout of OTX2 caused a predicted fate switch from rostral to caudal MHB (**Figure 6H**). In contrast, GBX2 was not identified as a key regulon in cMHB and its knockout concordantly did not cause any shift in MHB trajectories (**Figure 6H**), indicating that expression of GBX2 is not required for establishment of the human MHB. Knockout simulation of PAX2, PAX5 and EN1 in turn caused an instability of the early presumptive MHB populations, implicating these genes in early MHB specification, while knockout of PAX8 caused pronounced fate shifting from caudal to rostral MHB populations, thereby exhibiting the opposite effect of OTX2. Interestingly, while GBX2 appeared dispensible for cMHB specification, we identified MSX1 and MSX2 as key transcription factors in the cMHB population (**Figure 6G**). Aligned with this, simulated knockout of MSX1 led to disruption specifically of the cMHB population, inducing a strong predicted fate shift towards rMHB (**Figure 6H**).

### *OTX2*-*GBX2* interaction is not sufficient to induce MHB formation

Having outlined the temporal gene expression patterns and GRNs involved in the formation of the human MHB in vitro, we proceeded to deconstruct the MHB in vitro model to identify minimal requirements for MHB induction and maturation. In the first line of experiments, we investigated whether WNTa alone could serve as the primary driver of MHB fate. We differentiated H9 hESCs in a 2D monolayer with the WNT activator CHIR99021 (CHIR) at 0.8-1.0 μM from d0-9 (i.e. equivalent to the concentrations present in the C-region of the MiSTR). However, qRT-PCR analysis at day 14 of key MHB markers revealed negligible induction of MHB markers in this monolayer system compared to the MHB region in our MiSTR model, suggesting that WNTa alone is insufficient to induce MHB formation (**Figure 7A**). Next, we tested if the induction of MHB required WNTa in combination with 3D cellular interaction. For this, we differentiated cells in neural spheroid cultures which were exposed to discrete concentrations of CHIR spanning from diencephalic to hindbrain concentrations (0.4 to 1.4 μM). The qRT-PCR analysis at day 16 showed that while the 0.8 μM CHIR condition (= midbrain levels of WNT activation) induced upregulation of early MHB markers (*PAX5, PAX8, FGF8, EN1*), the expression of late MHB markers (*FGF17, WNT1* and *EN2*) remained low relative to that of MiSTR MHB region (**Figure 7A**). We next hypothesized that proper MHB induction and maturation might require additional juxtapositioning of OTX2-positive and GBX2-positive (OTX2-negative) cells, and thus created an OTX2^+^/GBX2^+^ interaction assay by fusing spheroids of OTX2^+^ midbrain (MB) and GBX2^+^ hindbrain (HB) identity. To generate MB spheroids, we used a CHIR concentration of 0.5 µM to avoid any hindbrain contamination, while 1.3 µM was used to generate HB spheroids devoid of midbrain contamination. We then fused MB and HB spheroids on day 6 (MB/HB fusion) and compared these to control fusions of MB-MB and HB-HB as well as to the MiSTR MHB region. In two of the fusion condition, we additionally added either WNTa (0.8 µM CHIR= midbrain levels) or FGF8 during the first three days post-fusion. Nonetheless, all fusion conditions failed to show significant induction of the mature MHB markers *FGF8, FGF17* and *EN2* in comparison to the MiSTR MHB region, indicating that OTX2-GBX2 interaction is not a key driver of MHB induction (**Figure 7A**).

**Figure 7.**
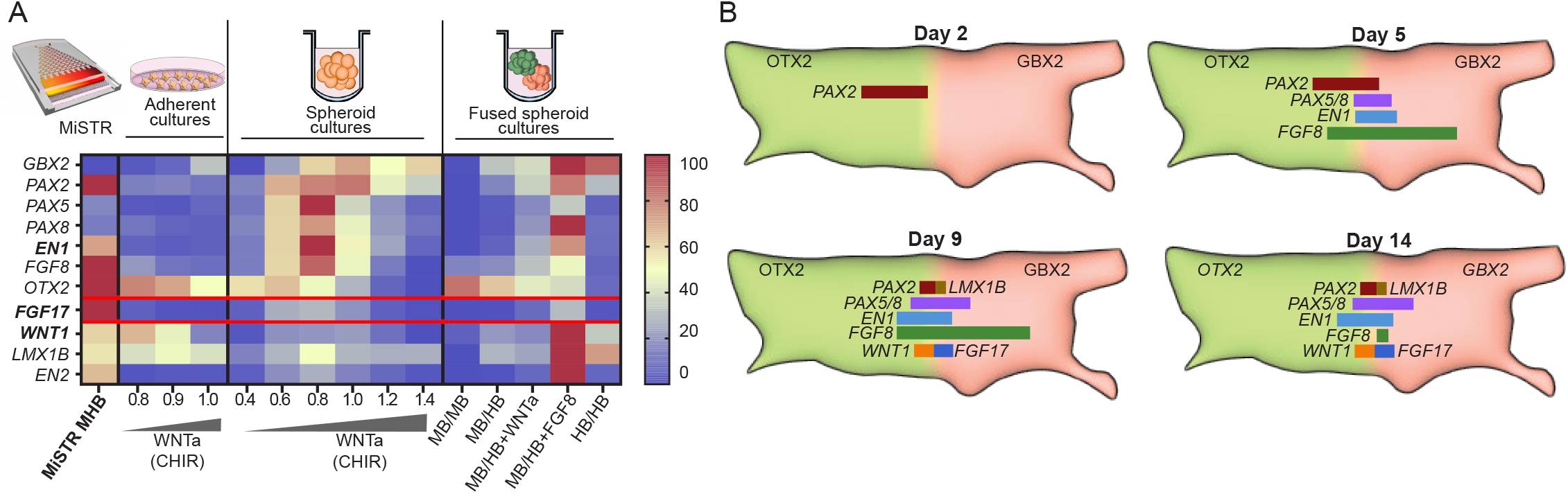
Mature MHB markers are not induced in alternative *in vitro* models. **A)** Comparative qRT-PCR analysis of MiSTR and three alternative culturing systems for testing induction of MHB fates (n = 3). Adherent monolayer cultures with different WNTa (CHIR) concentrations, spheroid cultures with different WNTa (CHIR) concentrations and fusion of midbrain (MB) and hindbrain (HB) spheroids at day 6 to model an OTX2/GBX2 interface in a 3D environment. **B)** Schematic summarizing the dynamics of MHB marker expression in MiSTR tissue as interpreted from scRNAseq and qRT-PCR data.

Together, these findings suggest that neither WNT activation alone nor OTX2*-*GBX2 interaction followed by WNT activation is sufficient to induce MHB formation. Instead, only the MiSTR model displayed efficient induction of mature MHB markers, indicating that early WNT signalling at exactly the right midbrain-level already at the epiblast stage is required for initiation of the MHB program. This will then induce early PAX2 expression in the caudal part of the OTX2+ domain, leading to subsequent activation of PAX5/8, EN1 and finally a clearly segregated expression of WNT1 and FGF17 in the OTX2^+^ rMHB and GBX2^+^ cMHB, respectively (**Figure 7B**). This data emphasises the importance of access to human in vitro models which can recapitulate complex aspects of neural tube patterning, including secondary signalling centers.

## Discussion

In recent years, significant advancements have been made in understanding embryonic and organ development through the utilization of scRNAseq atlases in various model organisms and humans. Notably, the scRNAseq atlases of mouse and human brain development, covering also early embryonic stages, have provided remarkable insights into neurodevelopment at the molecular level (Plass et al. 2018, Wagner et al. 2018, Pijuan-Sala et al. 2019, La Manno et al. 2021, Braun et al. 2022, Braun et al. 2023). However, investigating the early stages of human embryonic development within the 3-5 PCW timeframe, along the axis of these morphogenic gradients, poses significant challenges due to the limited availability of human fetal tissue, and the very small number of cells in the embryo at these early timepoints. To complement the fetal datasets at very early stages, we here analysed an extended MiSTR development dataset spanning the rostro-caudal and dorso-ventral axes. The MiSTR model allows us to deconstruct the developmental processes leading to neural tube regionalization in the absence of confounding extrinsic factors from neighbouring non-neural tissues in the embryo. Our findings demonstrate that by utilizing simple linear morphogenic gradients manipulating only two growth factor pathways – WNT and SHH – the MiSTR model can effectively recapitulate the main patterning features of early human neural tube development in the dish.

Taking advantage of having access to abundant cell numbers in the MiSTR model compared to the minute quantities available from early embryo tissues, we were able to uncover the earliest patterning events taking place in pre-neuralising cells. As has been indicated earlier (Rifes et al. 2020), we confirmed that rostro-caudal fate is established several days before the onset of neuralising transcription factors. This is in line with older data from the chick, showing that the anatomical features of neural plate invagination is established along the rostro-caudal axis of the head fold before the onset of *Pax6* expression in the neural plate region (Li et al. 1994). More recent studies in the early chick embryo have revealed that pluripotency genes, including Oct4 expression, remain active in the neural plate and neural fold regions of the chick up until neural tube closure at around stage 8/9 (Pajanoja et al. 2023). Transcriptional analyses further showed that the gene expression profile of the anatomical neural plate region during folding was remarkably similar to a pluripotent epiblast signature (Pajanoja et al. 2023). While the MiSTR lacks the embryonic anatomical features of the folding neural plate, we have in this study annotated these pre-neuralising cells as late rostral and caudal epiblast due to their high similarity to epiblast signatures. However, we believe these cells are equivalent to cells located in the folding neural plate of the embryo, before the onset of *SOX1* and *PAX6* expression. The high resolution of our MiSTR model across the rostro-caudal axis allowed us to identify the earliest rostral versus caudal priming factors expressed in the pre-neuralising cells. We identified *HESX1, SHISA2, FEZF1, LHX5* and *CYP26A1* as pre-neuralising factors specifically expressed in the rostral cells. These markers are likely to play an important role in inhibiting signalling from exogenous caudalising factors. Hesx1 is expressed in the most anterior head fold of the early E7.75 mouse embryo (Matsuda and Kondoh 2014) and has been shown to be required for forebrain development in zebrafish by directly inhibiting beta-catenin-mediated transcription (Andoniadou et al. 2011). Shisa2 resides in the endoplasmic reticulum where it inhibits maturation of Frizzled and FGF receptors, thereby cell-intrinsically inhibiting cellular responsiveness to FGFs and WNTs (Yamamoto et al. 2005). Early rostral expression of Shisa2 has also been found in the gastrulating chick embryo (Hedge and Mason 2008). In zebrafish, *Lhx5* has been shown to be essential for forebrain development by activating expression of the secreted Wnt antagonists *Sfrp1a* and *Sfrp5* (Peng and Westerfield 2006). Finally, CYP26A1 is one of the main enzymes responsible for degradation of retinoic acid (RA) – a morphogen with well-known roles in caudalisation of the hindbrain and spinal cord (Thatcher and Isoherranen 2009). Collectively, the pre-neuralising genes expressed in the rostral cells thereby likely serve to safeguard rostral fates by efficiently blocking caudalising signals from both WNTs, FGFs and RA. In contrast, pre-neuralised caudal cells express high levels of *CRABP2*; a transport protein which facilitates RA transfer and binding to its nuclear receptor complex (Pohl and Tomlinson 2020), thereby conversely enabling efficient responsiveness to RA. In line with the expression of these early fate safeguarding factors, the pre-neuralised rostral epiblast cells could not be converted to hindbrain cells by adding CHIR at day 2, and conversely, pre-neuralised caudal epiblast cells could not be reverted to forebrain by removing CHIR at day 2. From this data, we conclude that rostro-caudal fate specification takes place during a late epiblast-like transcriptional phase, before onset of neuralising transcription factors. Interestingly, the establishment of dorso-ventral axis in the telencephalon was in contrast a highly delayed event, becoming transcriptionally encoded only around 7-9 days of differentiation.

Subsequent to rostro-caudal and dorso-ventral regionalization, the tissues intrinsically initiates formation of secondary signalling centers as evident by highly cluster-specific growth factor secretion. We have previously reported the unique feature of the R/C MiSTR model in recapitulating MHB formation (Rifes et al. 2020). Here, we exploited this feature to deconstruct the factors required for establishment of a human MHB in vitro. It has been proposed that interactions between rostral *OTX2*^*+*^ cells and caudal *GBX2*^*+*^ cells play an important role in inducing MHB formation (Hidalgo-Sanchez et al. 2022). We found here that the juxtaposition of *OTX2*^*+*^ and *GBX2*^*+*^ cells, with or without simultaneous WNT activation, did not lead to proper induction of MHB markers. These results are somewhat inconsistent with previous observations made in chick/quail transplantations (Hidalgo-Sánchez et al. 1999) and explant fusions (Irving and Mason 1999). In line with this, *in silico* knockout studies on the MiSTR indicated that *GBX2* may not be essential for the formation of the MHB. This aligns with studies in *GBX2*^*-/-*^ mutant mice, which still show sustained formation of MHB structures with *Fgf8* and *Wnt1* expression (Millet et al. 1999). Instead, our MHB reconstruction studies suggest that a prerequisite for MHB induction is an early and precise priming of the late epiblast with midbrain-level WNT activation to initiate the MHB transcriptional program through early induction of *PAX2*. However, it is important to note that the later interaction between *OTX2*^*+*^ and *OTX2*^*-*^ cells does appear to be required for the induction of the late mature MHB markers *FGF17* and *EN2*, as these MHB markers are not induced in spheroids which have been exposed only to midbrain-level WNT activation.

Interestingly, *FGF17* was observed to have a broader, stronger and temporally more persistent expression profile than *FGF8* in the MiSTR MHB (**Figure S7)**. This is consistent with data from the human and mouse embryo, and prompted us to develop an alternative protocol using FGF17-based patterning to generate dopaminergic progenitors for cell replacement therapy in Parkinson’s disease (Nygaard et al. 2024). Additionally, we have also applied the MiSTR dataset to identify the ventral midbrain-specific cell surface marker *APCDD1* as a novel tool for quality control and cell purification for dopaminergic progenitor cell products for Parkinson’s disease (Kirkeby et al. 2023). These examples highlight the MiSTR scRNAseq dataset as a valuable resource for researchers to identify regional and subtype-specific marker expression profiles for improving translational stem cell therapy (Kirkeby et al. 2023). We have made the entire temporal MiSTR dataset easily accessible online as a resource to scientists interested in exploring early patterning events in human neural tube development.

## Data and Code availability

The processed datasets are available on https://cells.ucsc.edu/?ds=neural-tube-organoids. The data analysis notebooks are available at https://github.com/gauravsinghrathore/MiSTR_atlas. Raw sequencing data generated for this study will be deposited at GEO server and can be made available with a valid request.

## Author contributions

GSR, BT, JGC,THP and AK designed the study. GSR, FSC, MA, PR, CR. AHM. LSP,

UD, JK, and KLE performed experiments. GSR, EH, ZAN and JBC performed bioinformatics analysis and annotations. GSR and AK wrote the manuscript with input from all authors.

## Acknowledgements

We are grateful to the CBMR Single-Cell Omics Platform, J. Bulkescher from the reNEW Imaging Platform and Pablo Hernandez-Varas from the Core Facility for Integrated Microscopy (CFIM), University of Copenhagen for their assistance, technical expertise and facility access. We also thank Dr. Ilary Allodi and Dr. Roser Montañana-Rosell from Prof. Ole Kiehn Lab (Department of Neuroscience, University of Copenhagen) for assistance and support with RNAscope experiments. Special thanks to Martin Proks from reNEW for engaging scientific discussions and support with data analysis server. We would like to express our gratitude to Dr. Maximilian Haeussler and Dr. Marc Perry from UCSC. Genomics Institute for curating, hosting amd maintaining MiSTR dataset on UCSC cell browser. This study has been supported by funding from the Novo Nordisk Foundation (NNF18OC0030286, NNF17CC0027852 and NNF21CC0073729), the Lundbeck Foundation (R380-2021-1267 and R350-2020-963), Innovation Fund Denmark (BrainStem: 4108-00008A), EU H2020 (grant no, 874758) and the Knut and Alice Wallenberg Foundation. The funding bodies played no role in the design of the study and collection, analysis, and interpretation of data and in writing the manuscript.

## Methods

### hESC seeding for MiSTR

For the seeding of human embryonic stem cells (hESCs) in the MiSTR (Microfluidic Stem Cell Regionalisation) system, H9 hESCs (hPSCReg: WAe009-A, RRID: CVCL_9773, WiCell) were cultured in StemMACS iPS Brew XF medium (Miltenyi Biotec) on Matrigel-coated plates (20 µg/cm2, Corning). The cells were dissociated using UltraPure EDTA (0.5 mM, Invitrogen) and subsequently seeded onto a layer of Pure Phenol red free Matrigel matrix (Corning) bed placed on a polydimethylsiloxane (PDMS) bottom in iPS Brew medium with ROCK inhibitor. The assembled PDMS bottom with the seeded cells was then placed into the microfluidic MiSTR device and incubated overnight at 37°C.

### Assembly of MiSTR device

Assembly of the microfluidic cell culture system involved several steps to ensure sterility and proper functionality. Prior to each experiment, all components including the bottom and top PDMS modules, polycarbonate lid, tubing, and glass syringes were thoroughly cleaned with deionized water and autoclaved separately, except for the holding cassette and device holder which were sterilized using ethanol. The tubings connecting the glass syringes to the microfluidic gradient tree was checked for leaks and obstructions before each experiment.

Assembly took place inside a cell culture flow hood using sterilized MiSTR parts to maintain sterility and prevent clogging. On day 0, the top PDMS module and inlet tubing were wetted with sterile water and the water was immediately replaced by wash medium and bubble removal was ensured. This process was repeated with wash medium 2 hours later. Once the system was free of bubbles, the bottom PDMS module with the seeded cells was brought from the incubator, the temporary PDMS wall was removed, and the pluripotency medium was replaced with wash medium. The bubble-cleared top PDMS module was quickly assembled onto the bottom PDMS module, both submerged in wash medium to prevent bubble formation. The assembled device, along with the attached inlet tubing and media-filled glass syringes, was placed in the MiSTR holder and incubated at 37°C. To minimize pressure differences, the media-containing syringes were mounted on a syringe holder and connected to the neMESYS pumps. The flow of differentiation media was initiated, and the outlet tubing was connected to a sterile waste collector at a higher position to maintain a constant hydrostatic pressure and prevent bubble formation. The detailed methodology for assembling the microfluidic device for MiSTR tissue cuture in our previous MiSTR publication (Rifes et al. 2020).

### Differentiation to R/C MISTR tissue until day 14

The hESCs were differentiated in the MiSTR device using a continuous flow (160 μl/h) of neural progenitor medium (NPM). The NPM was composed of a 1:1 mixture of DMEM/F12 (50%, Gibco) and NeuroMedium (50%, Miltenyi Biotec), supplemented with N2 supplement (1:200, Gibco), NeuroBrew-21 without vitamin A (1:100, Miltenyi Biotec), Glutamax (1:200, Gibco), SB431542 (10 μM), and rh-Noggin (100 ng/ml). Two syringes were used, one containing medium with GSK3i (CHIR99021, Miltenyi Biotec) and the other without, the gradual mixing of both the media conditions in MISTR microfluidic device resulted in 0-100% of GSK3i gradient. To achieve ventralization, a combination of SHH-C24II (200 ng/ml) and purmorphamine (PurM) (0.5 μM) (both from Miltenyi Biotec) was added to the medium in both syringes. The syringes were refilled with fresh medium every 2-4 days. On day 9 of differentiation, the medium was changed to basal NPM without any added factors (SB+Noggin) or GSK3i gradient, and the flow rate was maintained until day 14.

### Differentiation to D/V MiSTR tissue until day 14

During the differentiation process in the MiSTR device, cells were cultured in neural progenitor medium (NPM) from day 0 to day 9. To establish a dorsal-ventral gradient, two conditions were prepared using the NPM medium supplemented with SB431542 and rh-Noggin. These conditions represented the 0% and 100% ends of the gradient. In the 100% condition, XAV939 (5 μM, STEMCELL Technologies) was added from day 0 to day 9, while SHH-C24II (200 ng/ml) and purmorphamine (0.3 μM) were added from day 3 to day 14 to promote ventralization. The 0% condition did not receive any additional factors to maintain dorsalization.

### Long term MiSTR culture (post day 14)

After day 14, the MiSTR tissue was removed from the MiSTR culturing microfluidic device. It was then horizontally divided into two equal halves (i.e. the tissue was cut in the middle along the long axis). The first half was further sectioned into five equal parts, which were dissolved in Qiagen RLT Plus buffer for subsequent quality control testing using qRT-PCR.

The second half of the tissue was utilized for ongoing culturing outside of the MiSTR device. To maintain the rectangular structure of the MiSTR tissue, it was embedded in matrigel and allowed to solidify at a temperature of 37 degrees Celsius. The matrigel-embedded tissue was then cultured in a maturation medium, which consisted of a mixture of DMEM/F12 (50%, Invitrogen) and MACS Neuro medium (50%, Miltenyi Biotec.). The medium was supplemented with Glutamax (1:200), Penicillin/Streptomycin (1:250) (Gibco/Invitrogen), B27 with vitamin A (1:100) (Gibco/Invitrogen), N2 supplement (1:200) (Gibco/Invitrogen), MEM-NEEA (1:200) (Gibco), insulin (1:40,000) (Sigma-millipore), and beta-ME (1:1000)(Gibco).

The embedded tissues were placed in flat-bottomed head culture plates placed on an orbital shaker. The culture medium was changed every 3-4 days to provide necessary nutrients and maintain optimal conditions for continued growth and development of the tissue.

### Single Cell/Nuclear seq of MiSTR Tissue

For the timepoints of day 0, 1, 2, and 14 of the R/C dorsal and ventral MiSTR tissues, previously sequenced libraries from (Rifes et al. 2020) were used. A similar approach to our prior MiSTR study (Rifes et al. 2020) was applied for the remaining MiSTR tissues to perform single cell RNA sequencing (scRNAseq).

For day 5, 9 (R/C dorsal, R/C ventral, and D/V forebrain), and day 14 (D/V forebrain) tissues, the tissues were dissected along the gradient. Half of each tissue was divided into five equal parts (regions A-E) for quality control using qRT-PCR mRNA analysis. The remaining half was also divided into five parts, dissociated into single cells with the Neural Dissociation Kit (P) from Miltenyi Biotec, and cryopreserved in CS10 CryoStor (Stemcell Technologies) at -150°C for future use. For days 21, 35 (R/C dorsal, R/C ventral, and D/V forebrain), and 62 MiSTR tissues (R/C dorsal, R/C ventral), which were in floating culture on shaker. The tissue was cut into 5 equal parts using spatula followed by similar cell dissociation were carried out using Neural Dissociation Kit (P) from Miltenyi, with 300,000 cells per part were reserved for qRT-PCR and the remaining stored at -150°C in CryoStor.

Following qRT-PCR quality control, cells from two to three replicates at each time point for each MiSTR model (R/C dorsal, R/C ventral, and D/V forebrain) and each part were thawed, pooled in 1.5 ml Eppendorf tubes in equal proportions in staining buffer (0.5% BSA in PBS), and incubated with unique cell hashing antihuman antibodies (five TotalSeq A0251–A0255 antibodies BioLegend) at 4°C for 30 minutes followed by 2X washes with staining buffer. The cells from all five regions were combined equally, and 25,000 cells were loaded onto a 10X Genomics lane using the V3.1 chemistry kit.

For day 62 MiSTR tissue, the dissociated cells underwent single nuclear RNA sequencing. The nuclear extraction was performed using the Nuclei Isolation Kit: Nuclei EZ Prep from (Sigma-Aldrich). Cells from all regions of the replicates were pooled in equal proportions and lysed using the EZ Lysis Buffer from the kit. The nuclei from the lysed cells were filtered using 50 µm filters.

The filtered nuclei were suspended in nuclei buffer containing PBS-/-with 1% BSA, 2 mM Mg2+, and 0.1% RNase inhibitor. They were then incubated with unique nuclear hashtag antibodies (five antibodies in total TotalSeq A0451–A455,BioLegend) for 30 minutes along with gentle flick in 15 minutes. The hashtagged nuclei from each region were resuspended in nuclei buffer with 0.5 µg/ml of DAPI, FACS sorted in equal numbers, and pooled. A total of 25,000 nuclei were loaded onto a 10X lane, and cDNA libraries were prepared using the 10X V3.1 kit.

### Regionalized neural monolayer differentiation

KOLF2.1J hiPSCs (hPSCreg: WTSIi018-B-12, RRID: CVCL_B5P3 (Pantazis et al. 2022) or H9 hESCs (hPSCReg: WAe009-A, RRID: CVCL_9773, WiCell) were cultured on 1 μg/cm^2^ Laminin-521 (Biolamina, #LN521-05) coated plates, maintained in StemMACS iPS Brew XF (Miltenyi Biotec, #130-530 104-368), and passaged when 70-90% confluent using 0.5 mM EDTA at 37°C (Thermo Fisher Scientific, #15575020). ROCK inhibitor (ROCKi; 10 μM; Y-27632, Miltenyi, #130-106-538) was added to the media for the first 24 hrs after passaging.

For monolayer differentiation into regionalized neural fates, the hPSCs were seeded on 2 μg/cm^2^ Laminin-521 coated plates in N2 media [50% MACS Neuromedium, 50% DMEM/F12 containing 1% N2 supplement (Thermo Fisher Scientific, #17502048), 1% GlutaMAX (Thermo Fisher Scientific, #35050061), and 10 U/mL Penicillin/Streptomycin (Gibco, #15070063)] including ROCKi for the first 24 hrs, and neuralisation was induced with 10 μM SB431542 (Miltenyi Biotech, #130-106-543), and 100 ng/mL Noggin (Miltenyi Biotec, #130-108-982) from day 0 to 9, as previously described (Nolbrant et al. 2017). For KOLF2.1 monolayer experiments, cells were differentiated towards four different regional fates through addition of different patterning factors: (1) dorsal forebrain (dFB): No added factors, (2) dorsal hindbrain (dHB): 2 mM CHIR99021 d0-9, (3) ventral diencephalon (vDienc): 300 ng/mL SHH-C24II (Miltenyi Biotec, # 130-095-730) d0-9, (4) ventral telencephalon (vTel): 5 mM XAV939 (STEMCELL Technologies) d0-9 and 300 ng/mL SHH-C24II d3-9. The medium was changed every second day, and RNA was collected on day 1, 2, 3, 4, 5, 7 and 9 after seeding. For H9 monolayer crossover experiments, cells were differentiated with either no added patterning factors (FB), or with 2 uM CHIR99021 added from d0-9 (HB), from d0-2 (HB->FB) or from d2-9 (FB->HB). All experiments were performed in biological triplicates, i.e. in three separate differentiation experiments.

### Adherent Culture Differentiations towards MHB Identity

H9 hESCs were differentiated towards MHB identity using a protocol modfied from (Nolbrant et al. 2017). On day 0, hESCs were seeded at a density of 10.000 cells/cm2 in NPM medium supplemented with SB431542 (SB, 10 μM, Miltenyi Biotec), rh-Noggin (Noggin, 100 ng ml/mL, Miltenyi Biotec) and 0.8 to 1.0 µM CHIR99021 (CHIR, Miltenyi Biotec) from day 0-9. From day 9-14, cells were cultured in NPM medium until day 14 at which point gene expression levels were analysed by RT-qPCR. Medium change was performed every second or third day following a 2-2-3 schedule with increasing amounts of medium.

### MHB Spheroid differentiations

H9 hESCs were differentiated towards neural progenitors as embryoid bodies (EBs) using a protocol modified from (Kirkeby et al, 2012). hESCs were seeded at a density of 350.000 cells/cm2 on non-treated multi-well plates (VWR) in NPM medium supplemented with SB431542 (SB, 10 μM, Miltenyi Biotec), rh-Noggin (Noggin, 100 ng ml/mL, Miltenyi Biotec) and CHIR99021 (CHIR, Miltenyi Biotec). Additionally, the medium contained Y-27632 (ROCKi, 10 µM, Miltenyi Biotec) from day 0 to day 2 to promote survival after dissociation. For CHIR titration experiment, Embryoid Bodies (EBs) were cultured in CHIR (0.4-1.4 µM) until day 9 and NPM until day 16. For spheroid fusion experiments, EBs were cultured in NPM supplemented with SB, Noggin and CHIR (0.5 µM for midbrain; 1.3 µM for hindbrain) until fusion on day 6. To fuse, two spheroids of approximately equal size and representing midbrain, hindbrain, or both were transferred to round-bottom, non-treated 96-wells on day 6. After fusion, EBs were cultured in NPM supplemented with SB and Noggin until day 9. Some cultures were additionally supplemented with CHIR (0.8 µM) or FGF8 (100 ng/mL, Miltenyi Biotec) from day 6 to 9. Hereafter, spheroids were cultured in NPM until day 16 at which point gene expression levels were analysed by RT-qPCR.

### qRT-PCR for hESCs, hiPSCs differentiation and MHB spheroids

For RNA collection, cells were lysed in RLT buffer (Qiagen, #74034) containing 0.5 mM beta-mercaptoethanol (Thermo Fisher Scientific, #31350010), snap frozen on dry ice, and stored at -80°C until RNA isolation. RNA was isolated with the QIAcube using the Rneasy Micro Plus kit (Qiagen, #74034) according to protocol, and cDNA was synthesized from 1 µg purified RNA using the Maxima First strand synthesis kit (Thermo Fisher Scientific, #K1642), diluted in EB buffer (Qiagen, #19086), and stored at -20°C until qRT-PCR analysis. For qRT-PCR, samples consisting of SYBR green (Roche, #04887352001), forward and reverse primer mix (see sequences in **Supplementary Table S4**), and cDNA were prepared in duplicates in a 384-well plate using a liquid dispensing robot iDOT (Dispendix), and the samples were run on the Light Cycler 480 II instrument (Roche, #05015243001). Ct values were measured over 40 cycles (60°C for 60 s annealing/elongation step and 95°C for 30s denaturation). The fold change of genes of interest was calculated as the average Ct value (average of the technical duplicates) normalized to the average expression of undifferentiated H9 and RC17 hESCs by the 2-ΔΔCt method (Livak and Schmittgen 2001), and then normalized to average expression of both *GAPDH* and *ACTB* housekeeping genes.

See **Supplementary Table S4** for a list of primers.

### Analysis of MiSTR MHB over time

qRT-PCR data from R/C dorsal MiSTR tissue regions A-E from day 0, 2, 5, 9, 14 was generated as described above. For each MiSTR, the MHB region was defined as the region (A-E) with the highest multiplication product of the OTX2 and GBX2 fold change expression. Most often the C-region contained the MHB. Subsequently, for each of the displayed genes, the mean fold change expression in the MHB region was normalized across day 0, 2, 5, 9 and 14 and visualized as a heatmap in GraphPad Prism 10.

### Statistics for qRT-PCR (MHB spheroids)

After computing the fold change in gene expression the statistical analysis was performed in GraphPad Prism 9. For all experiments, the alpha value was set at 0.05. For heat map visualisations, normalisation was performed using GraphPad, setting the lowest value of each gene to 0 and the highest value to 100. For comparing gene expression levels, one-way analysis of variance (ANOVA) test was used followed by Dunnett’s multiple comparisons test. All data points were tested for normal, or lognormal, distribution by the Shapiro-Wilk test in GraphPad. Datasets where (log)normality was not achieved were analysed using the one-way non-parametric ANOVA (Kruskal-Wallis) test using GraphPad.

### RNAscope

MiSTR tissue was sub-dissected to obtain one longitudinal (rostral to caudal) tissue-piece for RNAscope analysis and five tissue-pieces representing the A-E regions for qRT-PCR. Longitudinal tissue-pieces were embedded in OCT (Sakura) in Tissue-Tek cryomolds (Sakura), folded into a U-shape to minimise length and tissue fractures and frozen on an ethanol-dry ice mixture. OCT-embedded tissue samples were cryosectioned into 14-16 µm sections using a Cryostat Microm HM550 (Thermo) and mounted on Superfrost Ultra Plus slides (ThermoFisher Scientific). Sections were then stained using the RNAscope Multiplex Fluorescent Reagent Kit v2 Assay (ACD) with the following modifications: fixation in PFA 4% for 30 minutes and protease III (diluted 1:20 in PBS) treatment for 20 minutes. See **Supplementary Table S3** for a list of probes and fluorophores. Images were acquired through tile-scanning using either a Leica AF6000 fluorescence widefield screening microscope (20x, 0.4 NA) and LAS X software to perform tile-stitching, or the Axioscan 7 (Plan Apochromat 20x, 0.8 NA) and ZEN imaging software. Confocal imaging was performed using a Zeiss LSM 780 fluorescence confocal microscope and processed using FIJI (ImageJ, 1.53c).

### Immunostaining of MiSTR tissue

Immunostaining staining was performed on MiSTR tissue dissected, sectioned and mounted as described for RNAscope analysis. Tissue samples were fixated in PFA 4% for 30 minutes, washed twice in PBS and incubated for 1-3h in blocking buffer consisting of PBS, 0,1% Triton X-100 (Sigma) and 5% donkey serum (Biowest). Primary antibodies were diluted in blocking buffer and incubated overnight at 4ºC. Tissue samples were subsequently washed three times in PBS, incubated with secondary antibodies and DAPI for 2h at RT, and washed three times in PBS before imaging. See **Supplementary Table S2** for antibodies and dilutions.

### Whole mount immunocytochemistry

Whole MiSTR tissue samples were fixated in PFA 4% for 1 – 1.5 hours, washed in PBS, and permeabilized overnight in PBS supplemented with 0.2% Triton X-100 (Sigma-Aldrich), 20% DMSO (Fisher Scientific), and 0.3 M glycine (Sigma-Aldrich). Then the samples were incubated in blocking buffer consisting of 0.2% Triton X-100, 5% donkey serum, and 10% DMSO. Primary antibodies were diluted in PBS supplemented with 0.2% Tween-20 (Sigma-Aldrich), 0.01 mg/ml heparin (Sigma-Aldrich), 5% DMSO, and 3% donkey serum and samples were treated for 3 days. Tissues were extensively washed and incubated overnight in the solution consisting of secondary antibodies and DAPI diluted in PBS supplemented with 0.2% Tween-20 and 3% donkey serum and extensively washed in PBS. Αll the steps were performed at room temperature on an orbital shaker and all solutions were supplemented with 0.02% sodium azide. The day before imaging, samples were incubated in RapiClear 1.49 (Sunjin Lab) clearing solution. Samples were imaged on an inverted confocal microscope (ECLIPSE Ti2-E, Nikon) equipped with a spinning disk module (CSU-W1, Yokogawa) with 5x and 20x objectives. Images were processed in ImageJ (NIH).

### Sequence alignment and generating count matrices

The raw cDNA libraries from our previous MiSTR publication (Rifes et al. 2020) and newly sequenced cDNA libraries were processed and aligned against GRCh38-2020 human reference genome using 10X CellRanger version 5.0.1 and CellRanger version 3.1.0 for D/V forebrain MiSTR datasets to generate count martrices. The obtained CellRanger BAM files were aligned with the same reference genome using STARsolo version 2.7.10a (Kaminow et al. 2021) to generate intronic and exonic count matrices required for RNA velocity analysis. To ensure consistency of cells in both matrices, only cells with matching barcodes to the 10X CellRanger count matrices were selected. The hastag fastq files were processed using the CITE-seq-Count-1.4.3 (Stoeckius et al. 2017) to generate hashtag oligo (HTO) count matrices.

### Preprocessing, clustering, dataset integration

The resulting count matrices were initially processed with Seurat version 4.0.5 (Hao et al. 2021). HTO demultiplexing for days 5, 9, 14, 21, 35, and 62 samples was performed using the HTODemux Seurat function to assign each cell to its corresponding MiSTR tissue region. Cells identified as doublets were excluded from further analysis.To maintain data quality, cells exhibiting high mitochondrial content and gene counts surpassing twice the median genes per cell were identified as experimental outliers and excluded from the analysis. Furthermore, to mitigate potential clustering bias stemming from metabolism-related factors, mitochondrial and ribosomal genes were removed from further analysis. To mitigate the cell cycle effect and to reduce noise arising from unique molecular identifiers (UMIs), we applied SCTransform version 0.3.2 (Hafemeister and Satija 2019) to regress out the cell cycle score and RNA counts and compute variable features followed by Louvain clustering using Seurat function FindClusters. To further enhance the analysis, cluster markers enriched in each cluster were identified using the FindAllMarkers function in Seurat in order to enrich for top markers of each cluster. The MiSTR samples generated for this study and reanalysed data from our previous publication (Rifes et al. 2020) were integrated with FastMNN (batchelor, version 1.6.2) (Haghverdi et al., 2018) using RunFastMNN SeuratWrapper (version 0.3.2).

Potential neurons were identified based on high expression of the *STMN2*, and cells with an arbitrary expression level greater than 2.5 of the normalized *STMN2* expression value were subsetted into a new Seurat object. Subsequently, reclustering was performed on this subset using the Seurat function FindClusters. Similaraly FindAllMarkers function was used to identify top markers of each cluster.

The overall integrated MiSTR dataset from day 0-62 was then moved to SCANPY (Wolf et al. 2018) using SeuratDisk (Version: 0.0.0.9019, https://github.com/mojaveazure/seurat-disk). The data with counts was read in Scanpy preprocesed (normalized, log transform, scaled followed by identification of highly variable genes) using sc.pp.normalise_total, sc.pp.log1p and sc.pp.scale functions. The neighbors were identified using function sc.pp.neighbors based on MNN instead of PCA. Further leiden clustering was performed using sc.pl.leiden setting a resolution at 1.2 followed by enrichment of top cluster markers using sc.tl.rank_genes_groups based on Wilcoxon ranksum test (see code).

### Spatial Mapping

BoneFight (version 0.1.0) (La Manno et al. 2021) was used to project day 14 and 21 R/C dorsal, R/C ventral, and D/V forebrain MiSTR tissues toward publicly available post-conception week 5 human neural tube spatial data (https://storage.googleapis.com/linnarsson-lab-human/EEL_HE_5week/LBEXP20211113_EEL_HE_5w_970um_RNA_transformed_assigned.parquet) (Braun et al. 2023). The MiSTR dataset was subsetted accordingly. Additionally, prior to the alignment the following adjustments were made to improve its outcome. Firstly, we removed the region corresponding to the medullary hindbrain from the spatial data because it is not included in the MiSTR model. Furthermore, out of the genes shared with the spatial dataset, we identified the 25 most differentially expressed genes for each tissue from the subsetted MiSTR data, resulting in a total of 125 unique genes. The cluster level mean expression matrix was computed for these genes and projected against the subsetted spatial data using BoneFight. The results were locally smoothed (Braun et al., 2023), and the tissue with the highest probability value was assigned to each spatial spot, resulting in a spatial map of day 14 and 21 MiSTR tissue clusters. Tangram (version 1.0.3) (Biancalani et al. 2021) Python package was used to project the gene expression of these tissue clusters onto the spatial space.

### Comparative data analysis with the developing human brain atlas

The publicly available first-trimester developing human fetal brain scRNAseq dataset was downloaded (https://storage.googleapis.com/linnarsson-lab-human/HumanFetalBrainPool.h5) and read into anndata object. Since we were interested in early development, we subsetted timepoints from week 5.5. 6.6, 7.0, 7.5, 8.0. Additionally, for computational efficiency this dataset was randomly subsampled to 200K cells. We simplified the original tissue annotations by pooling them into Forebrain (including Cortex, Subcortical forebrain, Tel/diencephalon), Midbrain (including Mesencephalon, Thalamus, Hypothalamus, Ventral midbrain, Dorsal midbrain), and Hindbrain (including Cerebellum, Medulla, Pons, Pons/Cereb), this allowed an easier comparison with MiSTR tissue. To compare this dataset with MiSTR tissue, we selected MiSTR timepoints from day 9 to 62 with the same genes present in both datasets.

Standard scANVI (scvi-tools) (Lopez et al. 2018, Xu et al. 2021) workflow, was used to integrate the above described human fetal brain data into a reference. For this purpose, the scVI model was trained to remove the dataset-specific batch effect (batch_key: Age). The pre-trained scVI model was used to initialize the scANVI model which further corrects the batch effect, while optimizing the cell distances to ensure cells containing the same annotation (labels_key: Tissue) are closer together. After obtaining the developing human fetal brain reference dataset and its respective scANVI model, the model was used to predict tissues from the MiSTR dataset. This was achieved by retraining the reference scANVI model together with the MiSTR dataset. Using scANVI’s prediction function, the retrained model was applied to predict the unobserved cell types, which here corresponded to cells from the MiSTR dataset. The MDE and UMAP embeddings and Sankey plot were used for visualization. Pie charts were created using python librtary Matplotlib.

### Comparative data analysis with gastrulating mouse embryo

The publicly available gastrulating mouse embryo scRNAseq dataset was download from https://github.com/MarioniLab/EmbryoTimecourse2018/ to make it comparable with human it was mapped with human ortholog gene names. To study earliest neural patterning events we only selected cells from stage E7.0,E7.25,E7.5,E7.75 and E8.0 and subsetted these as a new object. Furthermore, to reduce model training complexity similar cell type annotations were merged as one for example Erythroid1, Erythroid2 and Erythroid3 were merged as Erythroid (see code). To compare this dataset with MiSTR tissue, we selected MiSTR timepoints from day 0 to 5 with the same genes present in both datasets. As previously described similar approach of implementing Standard scANVI (scvi-tools) (Lopez et al. 2018, Xu et al. 2021) workflow was used to integrate the gastrulating mouse dataset into a reference. For this purpose, mouse gastrulation dataset was used as reference and scVI model was trained to remove the dataset-specific batch effect (batch_key: stage). The pre-trained scVI model was used to initialize the scANVI model which further corrects the batch effect, while optimizing the cell distances to ensure cells containing the same annotation (labels_key: celltype) are closer together. After obtaining the reference dataset specific scANVI model, the model was used to predict celltypes from the MiSTR dataset. This was achieved by retraining the reference scANVI model together with the MiSTR dataset. Using scANVI’s prediction function, the retrained model was applied to predict the unobserved cell types, which here corresponded to cells from the MiSTR dataset. UMAP embeddings and Sankey plot were used for visualization.

### Visualization of Early MiSTR Patterning Data

To visualize the earliest genes expressed during rostro-caudal MiSTR patterning, cells from Day 0 (hESCs), Day 1, Day 2, and Day 5 of the R/C dorsal and R/C ventral MiSTR models were subsetted from the integrated dataset into a new AnnData object. The subsetted dataset was rescaled using the sc.pp.scale function in Scanpy. Principal component analysis (PCA) was performed using sc.tl.pca, followed by the computation of ForceAtlas2 (FA2) embeddings. Finally, the data was visualized with the sc.pl.draw_graph function.

### Monocle3 pseudotime inference of rostral and caudal fates

To study the transcriptional programs of early rostral (OTX2+) and caudal (GBX2+) patterning we performed a pseudotime analysis. For this purpose, we subtracted the day 0, 1, 2, 5, and 9 OTX2+ and GBX2+ cells into separate datasets using an expression cutoff determined with the Gaussian mixture model from scikit-learn (version 1.4.2), both the datasets were visualised on separate embeddings. Since day 0 cells were over-represented in the GBX2+ dataset, it was subsampled to 2500 cells using Scanpy. From both datasets, the top 2000 variable genes were detected followed by FastMNN batch integration. The pseudotime of caudal and rostral lineages was inferred using Monocle3 (version 1.3.7) (Cao et al. 2019). Gene expression of genes of interest was projected along pseudotime using polyfit function implemented in numpy (version 1.26.4).

### MiSTR early diencephalic development data analysis

The R/C ventral MiSTR tissue datasets from day 5, 9, 14, 21, and 35 were concatenated into a single dataset (adata object). We applied standard processing workflow using Scanpy where mitochondrial and ribosomal genes and cells expressing fewer than 200 genes or more than 2.5 times the median genes per cell were excluded. The resulting data was log-normalized followed by highly variable gene selection cell cycle gene regression and UMAP projection for data visualization. To identify cells expressing *NKX2-1* and *FOXA2*, we analyzed log-transformed gene expression values from the single-cell RNA-seq matrix. Density plots were generated for each gene to visualize their expression distributions. The cutoff for positivity was defined as the expression value where the first peak of the density curve began to settle, indicating the boundary between low and significant expression. Cells with expression values above this threshold were classified as positive for the respective gene (See code). The *NKX2-1* and *FOXA2* positive cells were subsetted into a new dataset where PCA was calucated using sc.tl.pca scanpy functions followed by computing nearest neighbors (sc.pp.neighnors) and further computing UMAP and ForceAtlas2 (sc.tl.umap and sc.tl.draw_graph) embedding. The adata was further cell cycle regressed and Louvain clustering (sc.tl.louvain) was used to compute clusters. Furthermore, wilcoxon rank sum tse was used to calculate cluster markers.

### MHB dataset analysis

To study the development dynamics of the Midbrain-Hindbrain Boundary (MHB) using the MiSTR model, we focused on the R/C dorsal MiSTR tissues from day 5, 9, and 14. We performed separate preprocessing (normalization, log transformation) for each time point followed by datasets concatenation and cell cycle regression. The final dataset integration was performed using the sc.tl.ingest function from scanpy(<u>tutorial</u>),. To assess MHB development, we calculated the MHB score for the integrated dataset using the sc.tl.score_genes function based on specific MHB genes (P*AX2, PAX8, FGF17, WNT1, FGF8, EN1*). Cells with a positive MHB score (>0.05) were extracted into a new anndata object, referred as MiSTR MHB dataset, for further analysis. In the MiSTR MHB dataset, we performed Louvain clustering at a resolution of 0.55 and computed UMAP and ForeceAtlas (FA2) embeddings for visualization. The MHB cells at this time points are highly dividing. Therefore, to look for differentially expressing transcription factors the cell cycle genes were deleated from the matirx followed by wilcoxon rank sum test using sc.tl.rank_genes_groups function. To elucidate the differentiation trajectories in MHB cells over time, we identified top 1500 highly variable genes and conducted RNA velocity analysis using the stochastic model from ScVelo Python library (Bergen et al. 2020). The resulting RNA velocity vectors were projected onto the FA2 embedding.

### GRN analysis and regulons enrichment

We utilized the Scenic protocol (Van de Sande et al. 2020) to identify regulatory factors of *NKX2-1+, FOXA2+*, and co-expressing cells. Here we used the OTX2+ expressing cells from the NKX2-1+/FOXA2+ dataset and performed regulon activity enrichment using GRNBoost2 and AUCell from pyScenic. Regulon specificity scores were computed for individual cell types (*NKX2-1+, FOXA2+*, and co-expressing cells) and timepoints (day 5, 9, and 14). To enrich regulons active in MHB development in the MiSTR tissue, we applied the Scenic protocol to the MHB dataset using GRNBoost2 and AUCell from pyScenic. Regulon specificity scores were calculated for the previously determined Louvain clusters, enabling us to identify regulons associated with MHB development.

### *In-silico* perturbation simulation

The in-silico perturbation of transcription factors was simulated using python library CellOracle (Kamimoto et al. 2023). For the NKX2-1+FOXA2+OTX2+ dataset, KNN imputation was performed with a K value of 162 and 50 principal components. The GRN analysis was performed using the transcription factor and target gene list obtained from SCENIC. Gene KO simulation was performed by setting the expression value of candidate genes to 0, and the resulting KO simulation vectors were projected onto the FA2 embedding, following the parameters provided in this tutorial from CellOracle authors: https://morris-lab.github.io/CellOracle.documentation/notebooks/05_simulation/Gata1_KO_simulation_with_Paul_etal_2015_data.html#Notebook-file.

For the simulation of gene Knock Out and overexpression in the MHB population, KNN imputation was performed with a K value of 293 and 50 principal components, and the GRN analysis was performed using the transcription factor and target gene list obtained from SCENIC. For gene KO simulation, the expression value of candidate genes was set to 0, while for overexpression, the double value of the highest expression of gene expression from the CellOracle object was selected. The resulting KO simulation vectors were visualized on the FA2 embeddings (see available code).

