## Supplementary Data for "Mapping early patterning events in human neural development using an *in-vitro* microfluidic stem cell model"

Figure S1

A

| Model | Time point | 10X Version | Est. no. of cells | Median genes per cell | Cells after filtering | HASHtagged in 5X regions | Biological replicates (n) | Cell ranger version |
|---|---|---|---|---|---|---|---|---|
| hESCs | Day 0 | V2 | 8333 | 2946 | 8197 | NO | NA | 5.0.1 |
| R/C Dorsal | Day 1 | V2 | 1667 | 2275 | 1033 | NO | 3 | 5.0.1 |
| | Day 2 | V2 | 4902 | 1647 | 4555 | NO | 3 | 5.0.1 |
| | Day 5 | V3 | 19632 | 2175 | 15965 | YES | 3 | 5.0.1 |
| | Day 9 | V3 | 16048 | 3530 | 13449 | YES | 3 | 5.0.1 |
| | Day 14 run 1 | V2 | 8091 | 2610 | 7597 | NO | 3 | 5.0.1 |
| | Day 14 run 2 | V2 | 5697 | 2736 | 5486 | NO | | 5.0.1 |
| | Day 14 run 3 | V2 | 6532 | 3183 | 6231 | NO | | 5.0.1 |
| | Day 21 | V3 | 17516 | 2565 | 15013 | YES | 3 | 5.0.1 |
| | Day 35 | V3 | 11671 | 2485 | 9145 | YES | 2 | 5.0.1 |
| | day 62 | V3 | 2689 | 1721 | 2564 | YES | 2 | 5.0.1 |
| R/C ventral | Day 1 | V2 | 9900 | 3183 | 8887 | NO | 3 | 5.0.1 |
| | Day 2 | V2 | 8576 | 3124 | 6547 | NO | 3 | 5.0.1 |
| | Day 5 | V3 | 16194 | 3897 | 9668 | YES | 3 | 5.0.1 |
| | Day 9 | V3 | 21458 | 2980 | 18024 | YES | 3 | 5.0.1 |
| | Day 14 | V3 | 16766 | 4064 | 13933 | YES | 3 | 5.0.1 |
| | Day 21 | V3 | 15437 | 2798 | 13249 | YES | 3 | 5.0.1 |
| | Day 35 | V3 | 9171 | 2038 | 7962 | YES | 2 | 5.0.1 |
| | day 62 | V3 | 6052 | 1968 | 5779 | YES | 2 | 5.0.1 |
| D/V forebrain | Day 5 | V3 | 13577 | 1951 | 7639 | YES | 2 | 3.1.0 |
| | Day 9 | V3 | 14907 | 1805 | 6437 | YES | 2 | 3.1.0 |
| | Day 14 | V3 | 14353 | 1984 | 6840 | YES | 2 | 3.1.0 |
| | Day 21 | V3 | 14099 | 1649 | 6223 | YES | 2 | 3.1.0 |
| | Day 35 | V3 | 14245 | 1552 | 5823 | YES | 2 | 3.1.0 |
| | | | Total 277513 | Average 2536 | Total 206246 | | Total tissue 54 | |

B

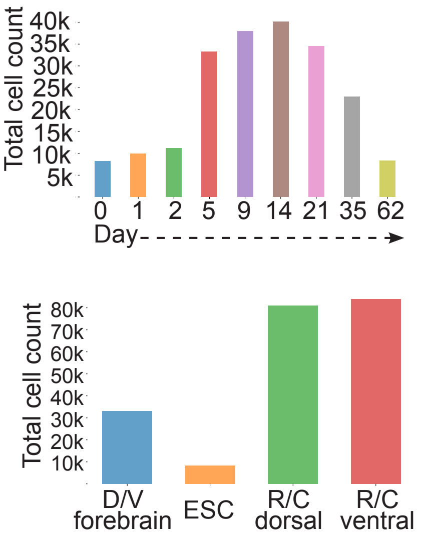


C

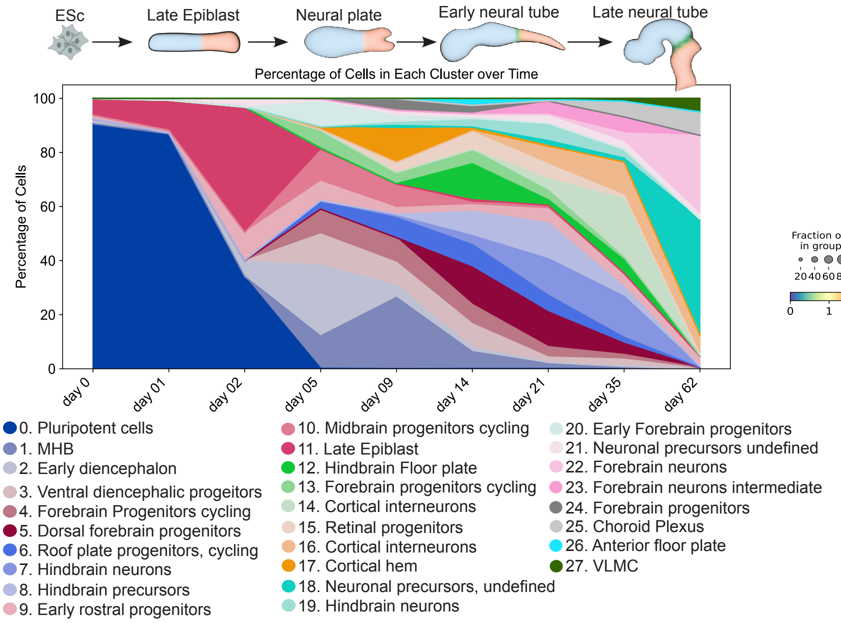


D

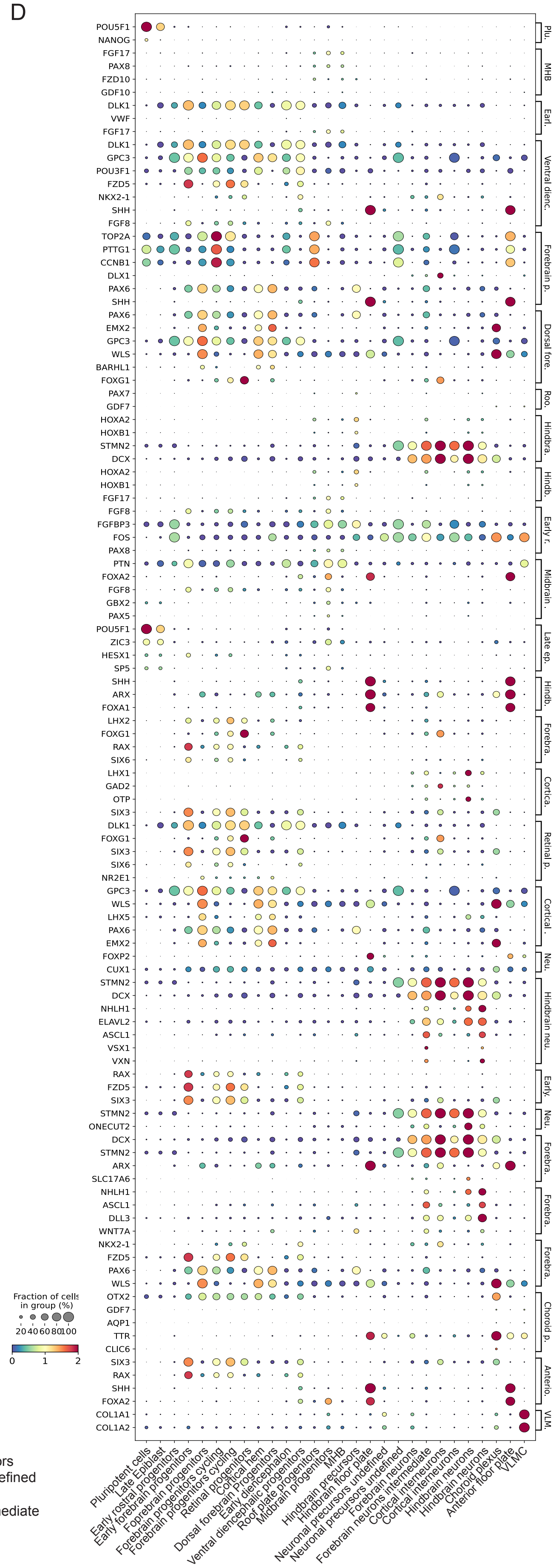

**Supplementary Figure S1. Marker gene expression in integrated MiSTR development dataset**

**A)** Table summarising 10X chromium experiment details. **B)** Complimenting histograms showing the number of cells sequenced form each day and model. **C)** Stackplot depicting the proportions of annotated clusters per timepoint showcasing a progressive increment in cellular heterogeneity throughout MiSTR development. The colors match with cell type colors from Figure 1. **D)** Dotplot showing average expression of key makers representing various celltypes in MiSTR dataset.

Figure S2

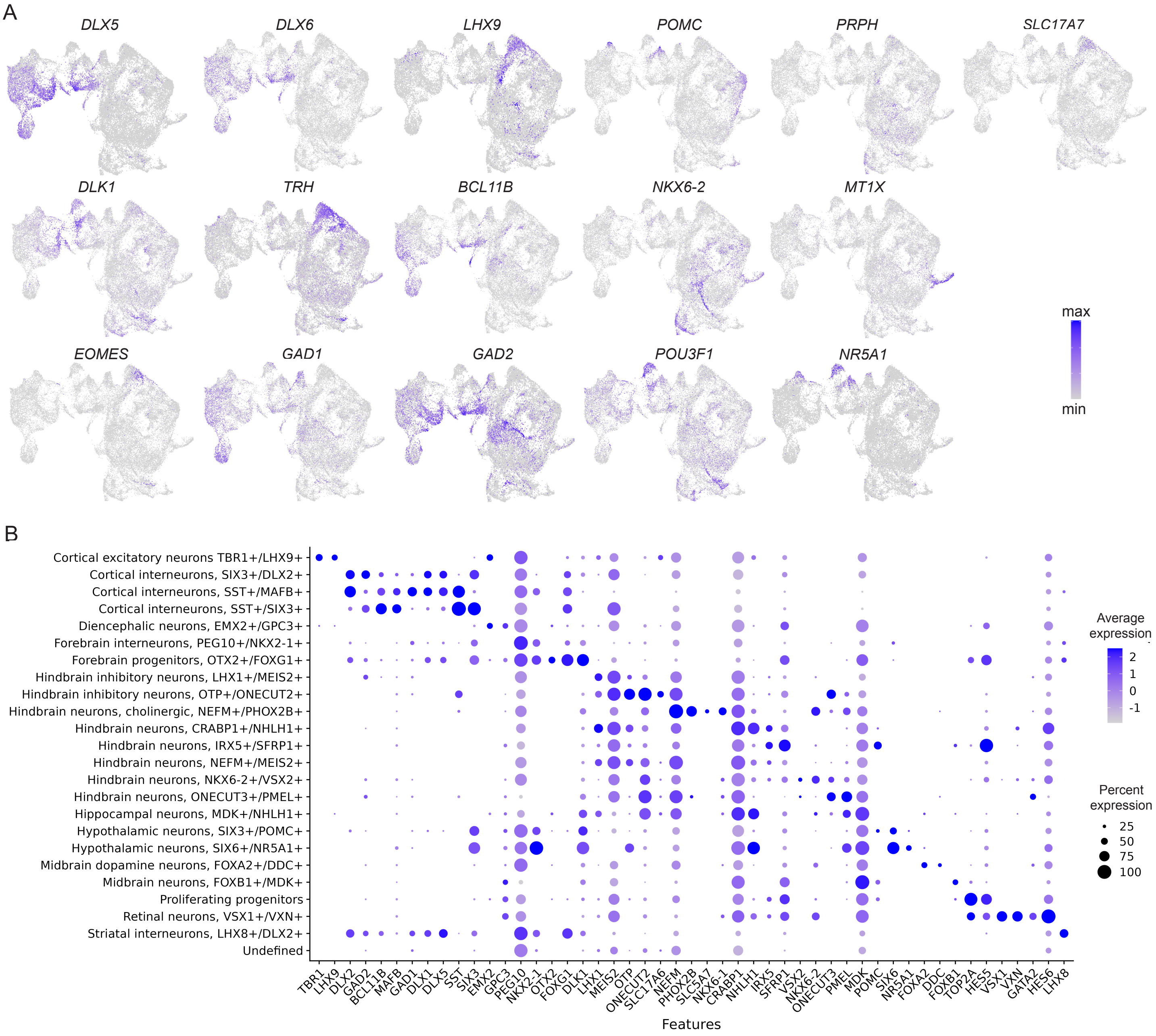
A
DLX5
DLX6
LHX9
POMC
PRPH
SLC17A7
DLK1
TRH
BCL11B
NKX6-2
MT1X
EOMES
GAD1
GAD2
POU3F1
NR5A1
max
min
B
Cortical excitatory neurons TBR1+/LHX9+
Cortical interneurons, SIX3+/DLX2+
Cortical interneurons, SST+/MAFB+
Cortical interneurons, SST+/SIX3+
Diencephalic neurons, EMX2+/GPC3+
Forebrain interneurons, PEG10+/NKX2-1+
Forebrain progenitors, OTX2+/FOXG1+
Hindbrain inhibitory neurons, LHX1+/MEIS2+
Hindbrain inhibitory neurons, OTP+/ONECUT2+
Hindbrain neurons, cholinergic, NEFM+/PHOX2B+
Hindbrain neurons, CRABP1+/NHLH1+
Hindbrain neurons, IRX5+/SFRP1+
Hindbrain neurons, NEFM+/MEIS2+
Hindbrain neurons, NKX6-2+/VSX2+
Hindbrain neurons, ONECUT3+/PMEL+
Hippocampal neurons, MDK+/NHLH1+
Hypothalamic neurons, SIX3+/POMC+
Hypothalamic neurons, SIX6+/NR5A1+
Midbrain dopamine neurons, FOXA2+/DDC+
Midbrain neurons, FOXB1+/MDK+
Proliferating progenitors
Retinal neurons, VSX1+/VXN+
Striatal interneurons, LHX8+/DLX2+
Undefined
TBR1
LHX9
DLX2
GAD2
BCL11B
MAFB
GAD1
DLX1
DLX5
SST
SIX3
EMX2
GPC3
PEG10
NKX2-1
OTX2
FOXG1
DLK1
LHX1
MEIS2
OTP
ONECUT2
SLC17A6
NEFM
PHOX2B
SLC5A7
NKX6-1
CRABP1
NHLH1
IRX5
SFRP1
VSX2
NKX6-2
ONECUT3
PMEL
MDK
POMC
SIX6
NR5A1
FOXA2
DDC
FOXB1
TOP2A
HES5
VSX1
VXN
GATA2
HES6
LHX8
Features
Average expression
2
1
0
-1
Percent expression
25
50
75
100

**Supplementary Figure S2. Marker gene expression in neuronal subset of integrated MiSTR dataset**

**A)** Gene expression plots of neuronal markers showing different subtypes of neurons.

**B)** Dot plot showing the top 5 cluster markers from the neuronal subset clusters from the integrated MiSTR dataset of all three models (day 0-62).

Figure S3

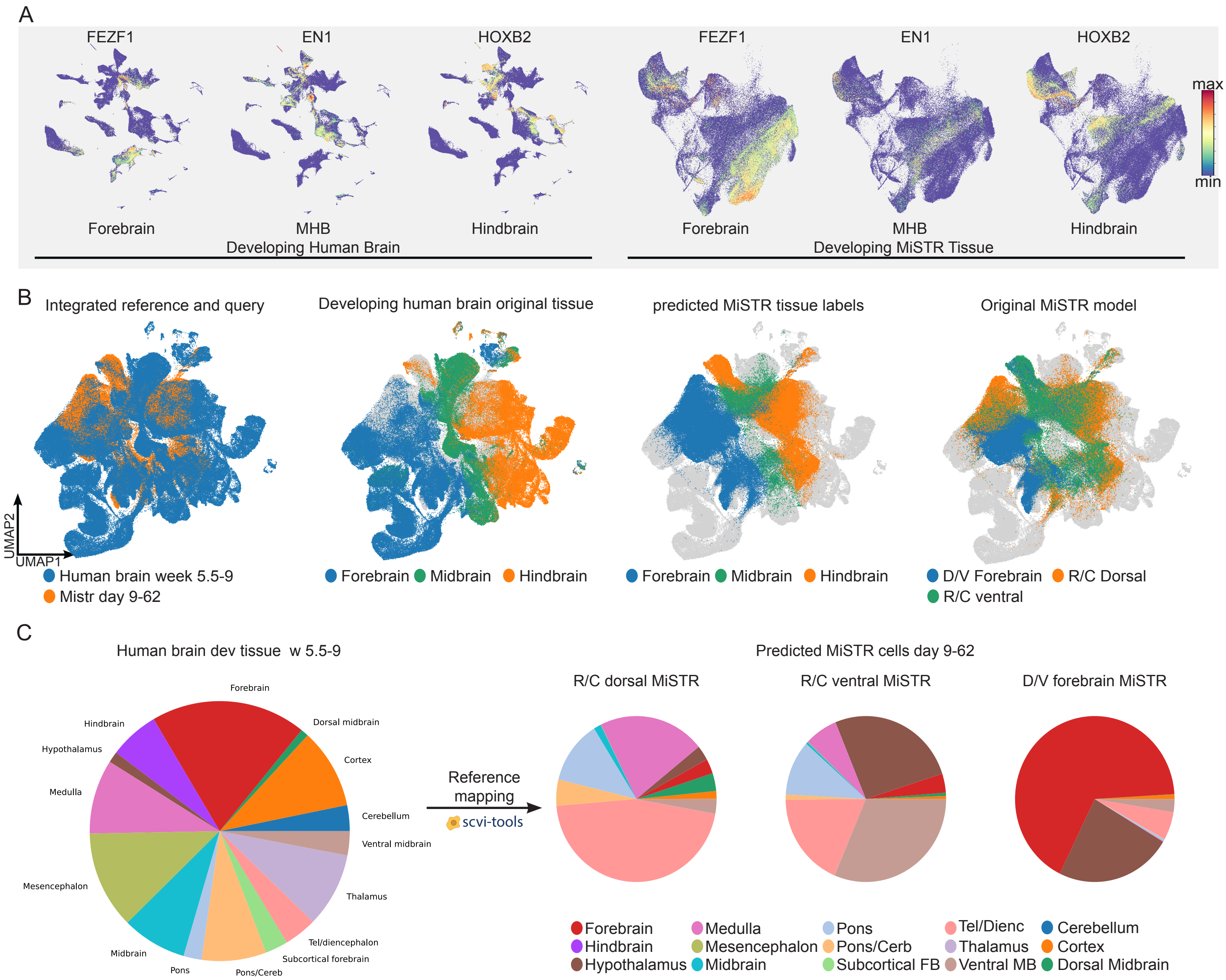
A
FEZF1
EN1
HOXB2
FEZF1
EN1
HOXB2
max
min
Forebrain
MHB
Hindbrain
Developing Human Brain
Forebrain
MHB
Hindbrain
Developing MiSTR Tissue
B
Integrated reference and query
Developing human brain original tissue
predicted MiSTR tissue labels
Original MiSTR model
UMAP2
UMAP1
Human brain week 5.5-9
Mistr day 9-62
Forebrain Midbrain Hindbrain
Forebrain Midbrain Hindbrain
D/V Forebrain R/C Dorsal
R/C ventral
C
Human brain dev tissue w 5.5-9
Forebrain
Hindbrain
Dorsal midbrain
Hypothalamus
Cortex
Medulla
Cerebellum
Ventral midbrain
Mesencephalon
Thalamus
Tel/diencephalon
Subcortical forebrain
Midbrain
Pons
Pons/Cereb
Reference
mapping
scvi-tools
Predicted MiSTR cells day 9-62
R/C dorsal MiSTR
R/C ventral MiSTR
D/V forebrain MiSTR
Forebrain
Hindbrain
Hypothalamus
Medulla
Mesencephalon
Midbrain
Pons
Pons/Cerb
Subcortical FB
Tel/Dienc
Thalamus
Ventral MB
Cerebellum
Cortex
Dorsal Midbrain

**Supplementary Figure S3. Mapping MiSTR tissue transcriptomes to the developing human brain atlas.**

**A)** Gene expression plots depicting side-by-side the region-specific markers *FEZF1* (Forebrain), *EN1* (midbrain-hindbrain boundary) and *HOXB2* (Hindbrain). **B)** UMAP plots of integrated reference (Braun et al*)* and query MiSTR dataset showing the efficieny of mapping where predicted cells in MiSTR tissue overlaps with the original query tissues. **C)** Pie chart showing proportion of cells from different brain tissues in developing human brain atlas followed by pie chart depicting the proportion of cells from the three MiSTR models (d9-62) mapped to different areas of the developing human brain atlas**.**

Figure S4

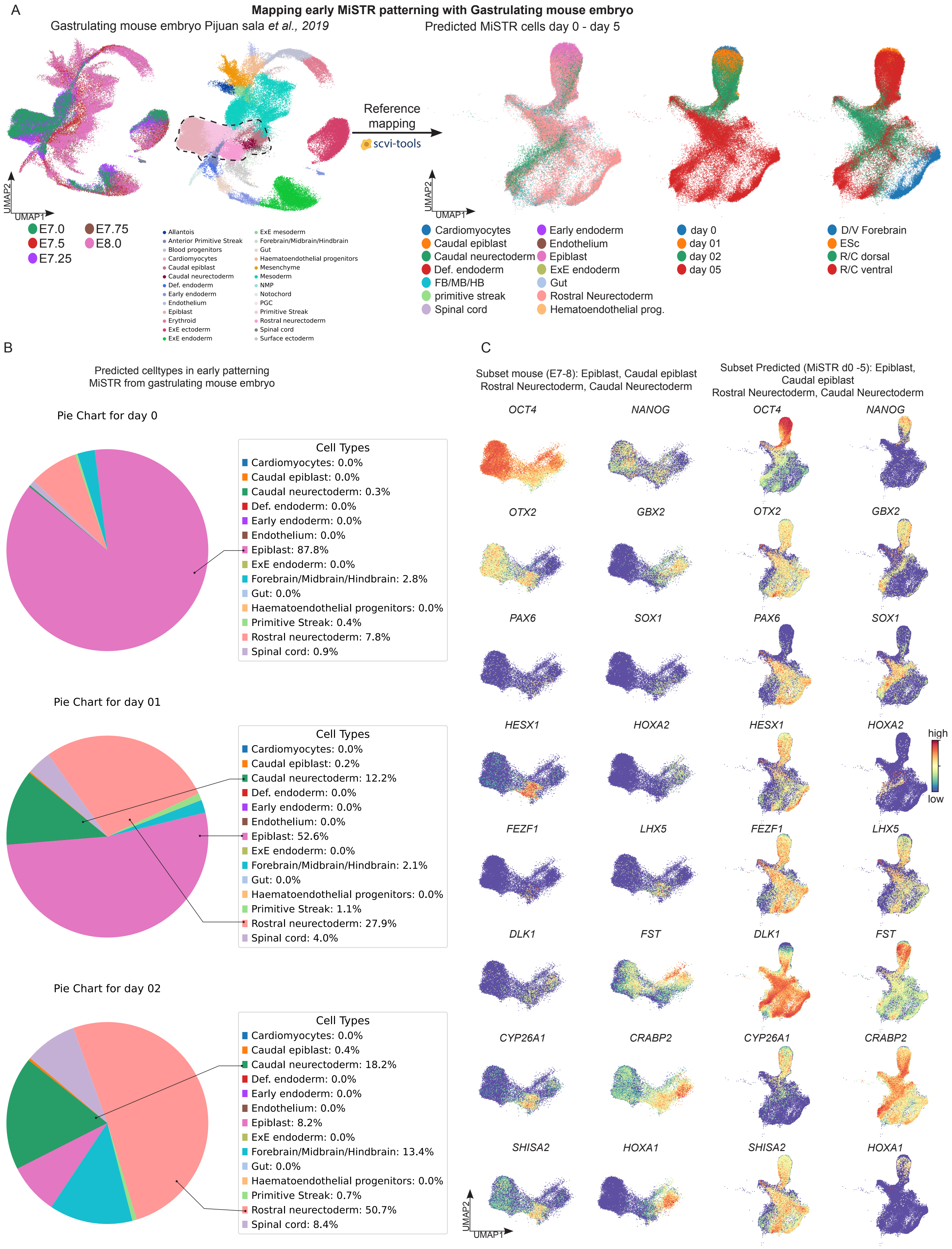
A
Mapping early MiSTR patterning with Gastrulating mouse embryo
Gastrulating mouse embryo Pijuan sala et al., 2019
Predicted MiSTR cells day 0 - day 5
Reference mapping
scvi-tools
UMAP2
UMAP1
E7.0
E7.5
E7.25
E7.75
E8.0
Allantois
Anterior Primitive Streak
Blood progenitors
Cardiomyocytes
Caudal epiblast
Caudal neurectoderm
Def. endoderm
Early endoderm
Endothelium
Epiblast
Erythroid
ExE ectoderm
ExE endoderm
ExE mesoderm
Forebrain/Midbrain/Hindbrain
Gut
Haematoendothelial progenitors
Mesenchyme
Mesoderm
NMP
Notochord
PGC
Primitive Streak
Rostral neurectoderm
Spinal cord
Surface ectoderm
Cardiomyocytes
Caudal epiblast
Caudal neurectoderm
Def. endoderm
FB/MB/HB
primitive streak
Spinal cord
Early endoderm
Endothelium
Epiblast
ExE endoderm
Gut
Rostral Neurectoderm
Hematoendothelial prog.
day 0
day 01
day 02
day 05
D/V Forebrain
ESc
R/C dorsal
R/C ventral
B
Predicted celltypes in early patterning MiSTR from gastrulating mouse embryo
Pie Chart for day 0
Cell Types
Cardiomyocytes: 0.0%
Caudal epiblast: 0.0%
Caudal neurectoderm: 0.3%
Def. endoderm: 0.0%
Early endoderm: 0.0%
Endothelium: 0.0%
Epiblast: 87.8%
ExE endoderm: 0.0%
Forebrain/Midbrain/Hindbrain: 2.8%
Gut: 0.0%
Haematoendothelial progenitors: 0.0%
Primitive Streak: 0.4%
Rostral neurectoderm: 7.8%
Spinal cord: 0.9%
Pie Chart for day 01
Cell Types
Cardiomyocytes: 0.0%
Caudal epiblast: 0.2%
Caudal neurectoderm: 12.2%
Def. endoderm: 0.0%
Early endoderm: 0.0%
Endothelium: 0.0%
Epiblast: 52.6%
ExE endoderm: 0.0%
Forebrain/Midbrain/Hindbrain: 2.1%
Gut: 0.0%
Haematoendothelial progenitors: 0.0%
Primitive Streak: 1.1%
Rostral neurectoderm: 27.9%
Spinal cord: 4.0%
Pie Chart for day 02
Cell Types
Cardiomyocytes: 0.0%
Caudal epiblast: 0.4%
Caudal neurectoderm: 18.2%
Def. endoderm: 0.0%
Early endoderm: 0.0%
Endothelium: 0.0%
Epiblast: 8.2%
ExE endoderm: 0.0%
Forebrain/Midbrain/Hindbrain: 13.4%
Gut: 0.0%
Haematoendothelial progenitors: 0.0%
Primitive Streak: 0.7%
Rostral neurectoderm: 50.7%
Spinal cord: 8.4%
C
Subset mouse (E7-8): Epiblast, Caudal epiblast Rostral Neurectoderm, Caudal Neurectoderm
Subset Predicted (MiSTR d0 -5): Epiblast, Caudal epiblast Rostral Neurectoderm, Caudal Neurectoderm
OCT4
NANOG
OTX2
GBX2
PAX6
SOX1
HESX1
HOXA2
FEZF1
LHX5
DLK1
FST
CYP26A1
CRABP2
SHISA2
HOXA1
high
low
UMAP2
UMAP1

**Supplementary Figure S4. Mapping MiSTR tissue transcriptomes to the gastrulating mouse atlas.**

**A)** Early timepoints of MiSTR tissue d0-5 mapped with gastrulating mouse embryo from E7 to E8 (Pijuan-Sala et al. 2019). **B)** Pie charts showing percentage of cells mapped with reference (mouse gastrulating atals) at d0 (hESC stage), d1 and 2 showing the pluripotent cells majorly mapped with epiblasts (pluripotent). Furthermore at d2 the percentage of predicted epiblasts increase along with rise of rostral and caudal neuroepithelium. **C)** UMAP plots of pluripotency marker *OCT4 (POU5F1),* along with rostral and caudal late epiblast patterning markers *OTX2, PAX6* and *GBX2, SOX1*.

Figure S5

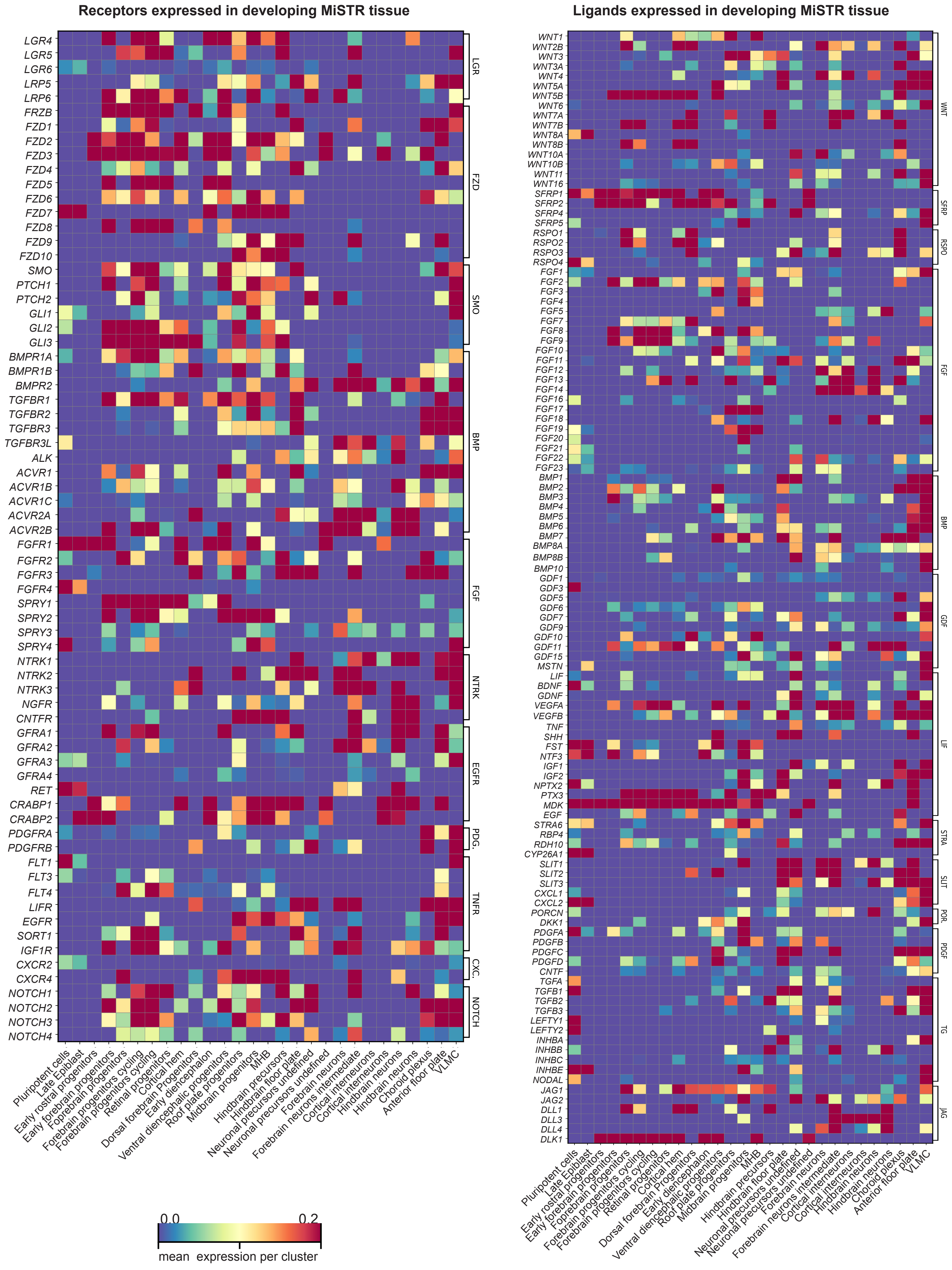
Receptors expressed in developing MiSTR tissue
Ligands expressed in developing MiSTR tissue
LGR
FZD
SMO
BMP
FGF
NTRK
EGFR
PDG.
TNFR
CXC.
NOTCH
WNT
SFRP
RSPO
GDF
LIF
STRA
SLIT
POR.
PDGF
TG
JAG
Pluripotent cells
Late Epiblast
Early rostral progenitors
Early forebrain progenitors
Forebrain progenitors
Forebrain progenitors cycling
Retinal progenitors
Cortical hem
Dorsal forebrain Progenitors
Early diencephalon
Ventral diencephalic progenitors
Roof plate progenitors
Midbrain progenitors
MHB
Hindbrain precursors
Hindbrain floor plate
Neuronal precursors undefined
Forebrain neurons
Forebrain neurons intermediate
Cortical interneurons
Hindbrain neurons
Choroid plexus
Anterior floor plate
VLMC
0.0
0.2
mean expression per cluster

**Supplementary Figure S5. Investigating secondary signalling centers in developing in all MiSTR models**

Matrix plot of secreted growth factors across all clusters of the integrated MiSTR dataset from all three models (d0-62), showing the emergence of region specific secondary signalling centers in the developing MiSTR tissues.

Figure S6

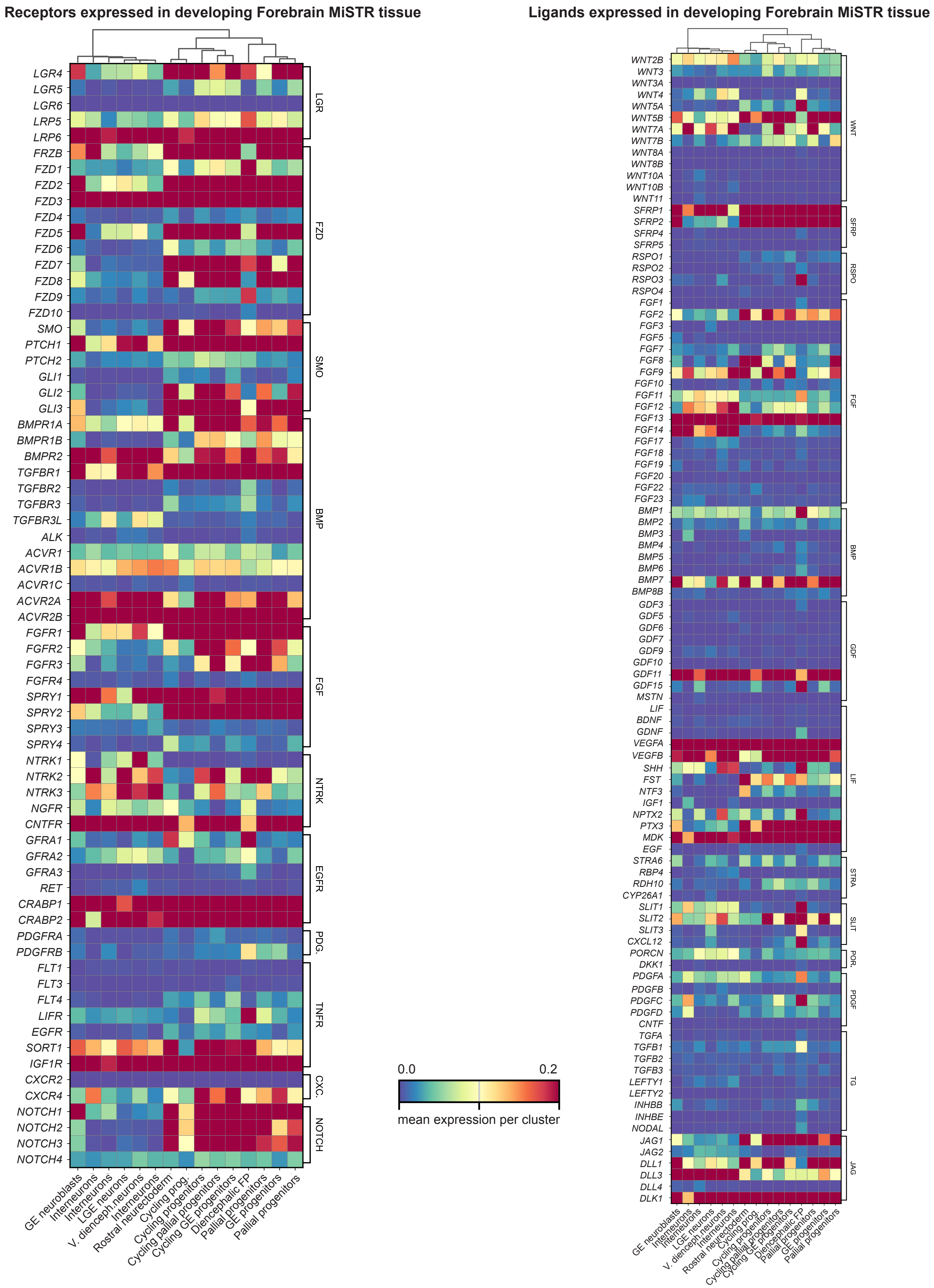
Receptors expressed in developing Forebrain MiSTR tissue
LGR4
LGR5
LGR6
LRP5
LRP6
FRZB
FZD1
FZD2
FZD3
FZD4
FZD5
FZD6
FZD7
FZD8
FZD9
FZD10
SMO
PTCH1
PTCH2
GLI1
GLI2
GLI3
BMPR1A
BMPR1B
BMPR2
TGFBR1
TGFBR2
TGFBR3
TGFBR3L
ALK
ACVR1
ACVR1B
ACVR1C
ACVR2A
ACVR2B
FGFR1
FGFR2
FGFR3
FGFR4
SPRY1
SPRY2
SPRY3
SPRY4
NTRK1
NTRK2
NTRK3
NGFR
CNTFR
GFRA1
GFRA2
GFRA3
RET
CRABP1
CRABP2
PDGFRA
PDGFRB
FLT1
FLT3
FLT4
LIFR
EGFR
SORT1
IGF1R
CXCR2
CXCR4
NOTCH1
NOTCH2
NOTCH3
NOTCH4
LGR
FZD
SMO
BMP
FGF
NTRK
EGFR
PDG.
TNFR
CXC.
NOTCH
GE neuroblasts
Interneurons
Interneurons
LGE neurons
V. dienceph.neurons
Interneurons
Rostral neurectoderm
Cycling prog.
Cycling progenitors
Cycling pallial progenitors
Cycling GE progenitors
Diencephalic FP
Pallial progenitors
GE progenitors
Pallial progenitors
0.0
0.2
mean expression per cluster
Ligands expressed in developing Forebrain MiSTR tissue
WNT2B
WNT3
WNT3A
WNT4
WNT5A
WNT5B
WNT7A
WNT7B
WNT8A
WNT8B
WNT10A
WNT10B
WNT11
SFRP1
SFRP2
SFRP4
SFRP5
RSPO1
RSPO2
RSPO3
RSPO4
FGF1
FGF2
FGF3
FGF5
FGF7
FGF8
FGF9
FGF10
FGF11
FGF12
FGF13
FGF14
FGF17
FGF18
FGF19
FGF20
FGF22
FGF23
BMP1
BMP2
BMP3
BMP4
BMP5
BMP6
BMP7
BMP8B
GDF3
GDF5
GDF6
GDF7
GDF9
GDF10
GDF11
GDF15
MSTN
LIF
BDNF
GDNF
VEGFA
VEGFB
SHH
FST
NTF3
IGF1
NPTX2
PTX3
MDK
EGF
STRA6
RBP4
RDH10
CYP26A1
SLIT1
SLIT2
SLIT3
CXCL12
PORCN
DKK1
PDGFA
PDGFB
PDGFC
PDGFD
CNTF
TGFA
TGFB1
TGFB2
TGFB3
LEFTY1
LEFTY2
INHBB
INHBE
NODAL
JAG1
JAG2
DLL1
DLL3
DLL4
DLK1
WNT
SFRP
RSPO
FGF
BMP
GDF
LIF
STRA
SLIT
POR.
PDGF
TG
JAG
GE neuroblasts
Interneurons
Interneurons
LGE neurons
V. dienceph.neurons
Interneurons
Rostral neurectoderm
Cycling prog.
Cycling progenitors
Cycling pallial progenitors
Cycling GE progenitors
Diencephalic FP
Pallial progenitors
GE progenitors
Pallial progenitors

**Supplementary Figure S6. Investigating secondary signalling centers in developing D/V forebrain MiSTR**

Matrix plot of secreted growth factors across all clusters of the integrated D/V Forebrain MiSTR dataset from d5 to 35, showing the emergence of region specific secondary signalling centers in the developing MiSTR tissues.

Figure S7

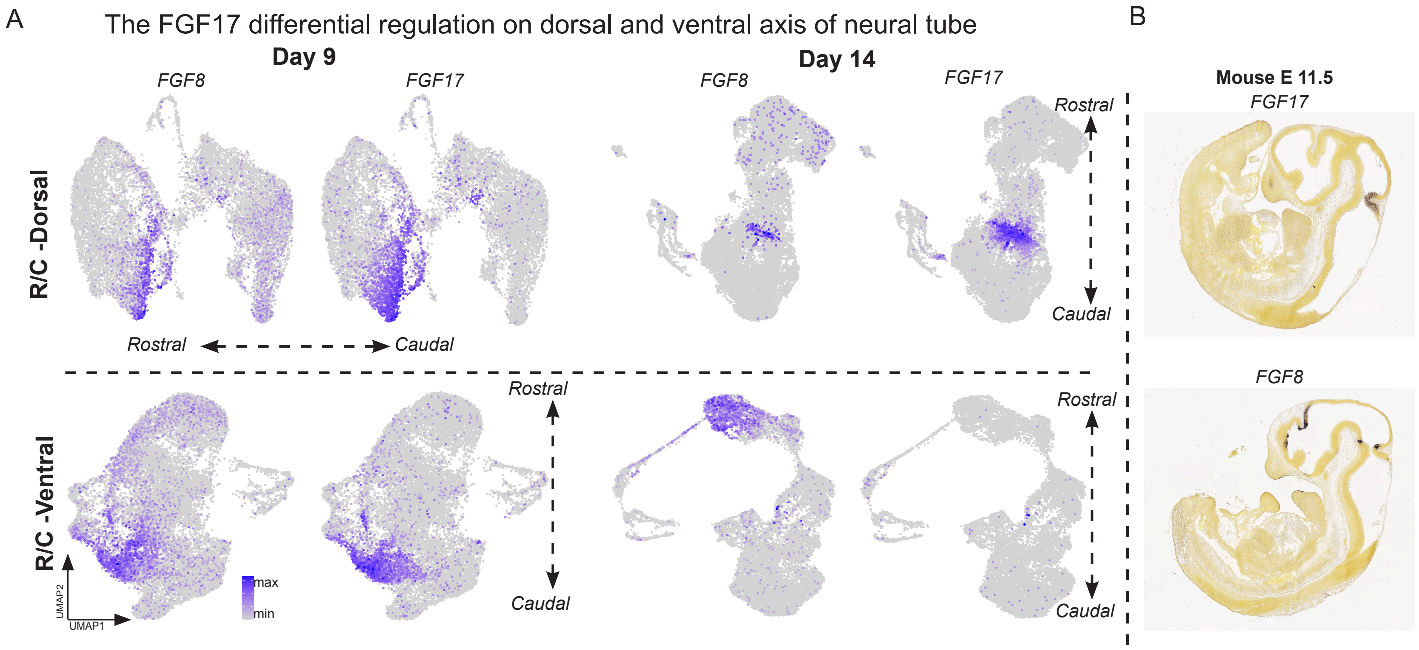
A
The FGF17 differential regulation on dorsal and ventral axis of neural tube
Day 9
Day 14
FGF8
FGF17
FGF8
FGF17
R/C -Dorsal
Rostral
Caudal
Rostral
Caudal
R/C -Ventral
Rostral
Caudal
Rostral
Caudal
UMAP2
UMAP1
max
min
B
Mouse E 11.5
FGF17
FGF8

**Supplementary Figure S7. Dorsal and ventral MHB dynamics and key regulons in MHB clusters**

**A)** *FGF8* and *FGF17* gene expression plots from R/C dorsal and R/C ventral MiSTR tissues (d9 and 14), depicting the temporal restriction of *FGF17* to the dorsal side of the MHB, whereas *FGF17* expression is lost in the ventral R/C MiSTR by day 14. **B)** This is similar to what is observed in the mammalian neural tube as visualised by in situ hybridisation images of *FGF8* and *FGF17* at E11.5 of the developing mouse embryo (Sourced from allen brain atlas).

### Supplementary tables

**Supplementary Table S1:**

Supplementary Table 1 is provided as a separate .xls file (see data availability).
This table contains cluster markers of Integrated MiSTR dataset along with subsetted neurons and individual MiSTR tissue datasets and their annotatios. Related to Figure 1.

**Supplementary Table S2:** Antibodies used for staining

| Primary antibodies | | | | | |
|---|---|---|---|---|---|
| **Epitope** | **Species** | **Source** | **Cat #** | **RRID** | **Dilution** |
| OTX2 | Goat | R&D Systems | AF1979 | AB_2157172 | 1:500 |
| FOXA2 | Mouse | Santa Cruz | sc-101060 | N/A | 1:500 |
| NKX2-1 | Rabbit | AbCam | ab133737 | N/A | 1:200 |
| SOX2 | Goat | R&D Systems | AF2018 | | 1:200 |
| FOXG1 | Rabbit | AbCam | ab18259 | AB_732415 | 1:500 |
| MAP2 | Mouse | Sigma-Aldrich | M1406 | | 1:500 |
| **Secondary antibodies** | | | | | |
| **Secondary Antibody** | **Host** | **Brand** | **Ref** | **RRID** | **Dilution** |
| Alexa Fluor 488 AffiniPure Donkey Anti-Goat IgG (H+L) | Donkey | Jackson ImmunoResearch | 705-545-147 | AB_2336933 | 1:200 |
| Cy™3 AffiniPure Donkey Anti-Rabbit IgG (H+L) | Donkey | Jackson ImmunoResearch | 711-165-152 | AB_2307443 | 1:200 |
| Alexa Fluor® 647 AffiniPure Donkey Anti-Mouse IgG (H+L) | Donkey | Jackson ImmunoResearch | 715-605-151 | AB_2340863 | 1:200 |
| 4',6-Diamidino-2-Phenylindole, Dilactate (DAPI) | – | Thermo | D3571 | N/A | 1 µg/mL |

**Supplementary Table S3:** Probes for RNAscope

| Probe Target | Channel | Brand | Ref |
|---|---|---|---|
| OTX2 | C3 | ACDBio | 484581-C3 |
| GBX2 | C1 | ACDBio | 483331 |
| PAX2 | C2 | ACDBio | 442541-C2 |
| EN1 | C2 | ACDBio | 527741-C2 |
| WNT1 | C3 | ACDBio | 429421-C3 |
| FGF8 | C2 | ACDBio | 415791-C2 |
| FGF17 | C1 | ACDBio | 1148351-C1 |

| Fluorophore | Brand | Ref |
|---|---|---|
| Opal 520 | Opal | FP1487001KT |
| Opal 570 | Opal | FP1488001KT |
| Opal 690 | Opal | FP1497001KT |

**Supplementary Table S4:** Human primer sequence for qRT-PCR

| Primer | Gene (Full Name) | FWD sequence | REV sequence |
|---|---|---|---|
| **ACTB** | Actin beta | ATGTGGCCGAGGACTTTGATTG | ATGGCAAGGGACTTCCTGTAAC |
| **ARX** | aristaless related homeobox | CCTGAGCACTTTCCTCGGAGCG | TGGAAAAGAGCCTGCCGAATGCC |
| ***CYP26A1*** | Cytochrome P450 Family 26 Subfamily A Member 1 | CCTTAGGAGCTGTGTAGGCAAA | CTGGCCAGCTCCACTGTAAATA |
| **BARHL1** | BarH like homeobox 1 | GTACCAGAACCGCAGGACTAAA | AGAAATAAGGCGACGGGAACAT |
| ***CRABP2*** | Cellular Retinoic Acid Binding Protein 2 | TTGCAGGGTCTTGCTTTCTTTG | CAGCCATTCCTCTTTGTTGGTG |
| **CORIN** | corin, serine peptidase | CATATCTCCATCGCCTCAGTTG | GGCAGGAGTCCATGACTGT |
| **DCX** | doublecortin | AGAAGCCATCAAACTGGAGACC | TCAGGACCACAGGCAATAAACA |
| **DLK1** | delta like non-canonical Notch ligand 1 | CTTTCGGCCACAGCACCTAT | CCTGCAAACATTGTCATCCTCG |
| **DLK2** | distal-less homeobox 2 | ACCAGACCTCGGGATCCGCC | CTGCGGGGTCTGAGTGGGGT |
| **EMX2** | empty spiracles homeobox 2 | AATCCACGCCTGCAAATTCTTC | TCCCGAGAATCTTGCACCAAA |
| **EN1** | engrailed homeobox 1 | CGTGGCTTACTCCCCATTTA | TCTCGCTGTCTCTCCCTCTC |
| **EN2** | engrailed homeobox 2 | CCTCCTGCTCCTCCTTTCTT | GACGCAGACGATGTATGCAC |
| **FEZF1** | FEZ family zinc finger 1 | GGTACATTCCACATTCGTGAGC | TCACGTGCAATAATCAAAACCA |
| **FGF17** | Fibroblast growth factor 17 | AAATCTGCTTCTCGGATCTCCC | CTACAGTCTAGCCAGGAGGAGT |
| **FGF8** | fibroblast growth factor 8 | ACAGCGCTGCAGAATGCCAAGT | GAAGTGGACCTCACGCTGGTGC |
| **FOXA1** | forkhead box A1 | GGGCAGGGTGGCTCCAGGAT | TGCTGACCGGGACGGAGGAG |
| **FOXA2** | forkhead box A2 | CCGTTCTCCATCAACAACCT | GGGGTAGTGCATCACCTGTT |
| **FOXG1** | forkhead box G1 | TGGCCCATGTCGCCCTTCCT | GCCGACGTGGTGCCGTTGTA |
| ***FST*** | Follistatin | GATGGGAAAACCTACCGCAATG | CTGGCTGCTCTTTACATCTTGC |
| **FZD5** | frizzled class receptor 5 | GCCTGTCGCTAAACTTTCCG | CGAGAAGAGCAGACAGTCCC |
| **GAPDH** | glyceraldehyde-3-phosphate dehydrogenase | TTGAGGTCAATGAAGGGGTC | GAAGGTGAAGGTCGGAGTCA |
| **GBX2** | gastrulation brain homeobox 2 | GTTCCCGCCGTCGCTGATGAT | GCCGGTGTAGACGAAATGGCCG |
| **GDF7** | growth differentiation factor 7 | GACGCTGCTCAACTCCATGGCA | TTGGCGGCGTCGATGTAGAGGA |
| **GLI1** | GLI family zinc finger 1 | CACTGTCCATCTCACCTCTGTC | GCGAGTTGATGAAAGCTACGAG |
| **GLI2** | GLI family zinc finger 2 | CAACCCTGTCGCCATTCACAAG | CACCCACATGAGCCGTGTC |
| **GLI3** | GLI family zinc finger 3 | GGGCCATCCACATGGAATATCT | CGGAGAGTGGTGATATGGACAG |
| ***HESX1*** | HESX Homebox 1 | TCACCCAATGCCAGAAGAAAGA | CTTGGTCTTCGGCCTCTATACC |
| **HOXA1** | homeobox A1 | GTACGGCTACCTGGGTCAAC | ACTTGGGTCTCGTTGAGCTG |
| **HOXA2** | homeobox A2 | CGTCGCTCGCTGAGTGCCTG | TGTCGAGTGTGAAAGCGTCGAGG |
| **HOXA3** | homeobox A3 | GGCCAATCTGCTGAACCTCA | GAGTTCAGATAGCCACCGGC |
| **HOXA5** | homeobox A5 | TCATAGTTCCGTGAGCGAGC | ATCCATGCCATTGTAGCCGT |
| **HOXB1** | homeobox B1 | GGCCTTCTCAGTACTACCCTCT | CCGTAGCTCGAGGGATGAAAAT |

| | | | |
|---|---|---|---|
| **IRX1** | Iroquois homeobox 1 | ACAGCAGTTAAAGTCGCCCTTC | AAGAAGACCCTTAATCAGGCGG |
| **IRX3** | iroquois homeobox 3 | GGCTTGCGCCCCGTAGAAATGT | AGGAGCCAGGTCAGGTCCGAAC |
| **LHX2** | LIM homeobox 2 | GGGCGACCACTTCGGCATGAA | CGTCGGCATGGTTGAAGTGTGC |
| **LHX2** | LIM homeobox 2 | GGGCGACCACTTCGGCATGAA | CGTCGGCATGGTTGAAGTGTGC |
| **LMX1A** | LIM homeobox transcription factor 1 alpha | CGCATCGTTTCTTCTCCTCT | CAGACAGACTTGGGGCTCAC |
| **LMX1B** | LIM homeobox transcription factor 1 beta | CTTAACCAGCCTCAGCGACT | TCAGGAGGCGAAGTAGGAAC |
| ***NANOG*** | Nanog Homebox | TTGGGACTGGTGGAAGAATC | GATTTGTGGGCCTGAAGAAA |
| **NKX2-1** | NK2 homeobox 1 | AGGGCGGGGCACAGATTGGA | GCTGGCAGAGTGTGCCCAGA |
| **NKX2-2** | NK2 homeobox 2 | CCCTGCTTTCGTGCATTTTGTA | ACTTGCTTTAGAAGACGGCTGA |
| **NKX6-1** | NK6 homeobox 1 | GGATCCCAACTCGGACGACGAGA | AGGATGAGCTCTCCGGCTCGG |
| ***LHX5*** | LIM homeobox 5 | TTGGACGAGTCTGAGATGTTGG | CGTAGTAGTCGCCTTGGTAGTC |
| **OLIG2** | oligodendrocyte transcription factor 2 | AAATCGCATCCAGATTTTCGGG | GAAGATAGTCGTCGCAGCTTTC |
| **OTP** | orthopedia homeobox | GAGTCCCGAGTGCAGGTCTGGT | GCACGGAACACGTTGGTCGTCT |
| **OTX2** | orthodenticle homeobox 2 | ACAAGTGGCCAATTCACTCC | GAGGTGGACAAGGGATCTGA |
| **PAX2** | paired box 2 | CTTGATGCCCCCTCCCCCGA | CCACTTCCACCCCGCAGCAG |
| **PAX5** | paired box 5 | CCCCATTGTGACAGGCCGTGAC | TCAGCGTCGGTGCTGAGTAGCT |
| **PAX6** | paired box 6 | TGGTATTCTCTCCCCCTCCT | TAAGGATGTTGAACGGGCAG |
| **PAX6** | paired box 6 | TGGTATTCTCTCCCCCTCCT | TAAGGATGTTGAACGGGCAG |
| **PAX8** | paired box 8 | ATAGCTGCCGACTAAGCATTGA | ATCCGTGCGAAGGTGCTTT |
| **POU3F1** | OCT6 POU class 3 homeobox 1 | GGACTCTTTTTGTTCGGTTGCT | TCGTCCGGGTGTTTGGTTTT |
| **POU5F1** | OCT4, POU class 5 homeobox 1 | TCTCCAGGTTGCCTCTCACT | GTGGAGGAAGCTGACAACAA |
| **PTCH1** | patched 1 | CATTGTACCTCGGGAAACCAGA | GCTGGATATTCGGGTAGTCTGC |
| **RAX** | Retina and anterior neural fold homebox | CCTCTCAGTTCACCAAGCAGAT | TGATCAACCTTGGGTGTTAGGG |
| **SHH** | sonic hedgehog | CCAATTACAACCCCGACATC | AGTTTCACTCCTGGCCACTG |
| ***SHISA2*** | Shisa Family member 2 | GCATCCAAGGTTAAGGGGAAGA | TGTGCCTGTGGAATACTGAGAC |
| **SIX3** | SIX homeobox 3 | ACCGGCCTCACTCCCACACA | CGCTCGGTCCAATGGCCTGG |
| **SIX6** | SIX homeobox 6 | CTCAACAAGAATGAGTCGGTGC | ACTCCTTGGTGAACTTGTGGTT |
| ***SOX1*** | SRY-box 1 | GGGAAAACGGGCAAAATAAT | TTTTGCGTTCACATCGGTTA |
| **SOX10** | SRY-box 10 | CTTTCTTGTGCTGCATACGG | AGCTCAGCAAGACGCTGG |
| **SOX9** | SRY-box 9 | CCACCCGGATTACAAGTACCAG | GAAGATGGCGTTGGGGGAGAT |
| **TBR1** | T-box, brain 1 | TCGTCCCCGCTCAAGAGCGA | CCTTGGCGCAGTTCTTCTCGCA |
| **TCF7L2** | transcription factor 7 like 2 | TTCCCTCCCCATATGGTCCC | TGTTGGTGTGACTATGGCCG |
| **WNT1** | Wnt family member 1 | GAGCCACGAGTTTGGATGTT | TGCAGGGAGAAAGGAGAGAA |
| **WNT3A** | Wnt family member 3A | GCGATGGCCCCACTCGGATACT | TAGCTGCCCAGAGCCTGCTTCA |
